# A Synthetic Cytotoxic T cell Platform for Rapidly Prototyping TCR Function

**DOI:** 10.1101/2023.11.20.567960

**Authors:** Govinda Sharma, James Round, Fei Teng, Zahra Ali, Chris May, Eric Yung, Robert A. Holt

## Abstract

Current tools for functionally profiling T cell receptors with respect to cytotoxic potency and cross-reactivity are hampered by difficulties in establishing model systems to test these proteins in the contexts of different HLA alleles and against broad arrays of potential antigens. We have implemented and validated a granzyme-activatable sensor of T cell cytotoxicity in a novel universal prototyping platform which enables facile recombinant expression of any combination of TCR-, peptide-, and class I MHC-coding sequences and direct assessment of resultant responses. This system consists of an engineered cell platform based on the immortalized natural killer cell line, YT-Indy, and the MHC-null antigen-presenting cell line, K562. These cells were engineered using contemporary gene-editing techniques to furnish the YT-Indy/K562 pair with appropriate protein domains required for recombinant TCR expression and function in a non-T cell chassis, integrate a fluorescence-based target-centric early detection reporter of cytotoxic function, and deploy a set of protective genetic interventions designed to preserve antigen-presenting cells for subsequent capture and downstream characterization. Our data show successful reconstitution of the surface TCR complex in the YT-Indy cell line at biologically relevant levels. We also demonstrate successful induction and highly sensitive detection of antigen-specific response in multiple distinct model TCRs, with significant responses (p < 0.05 and Cohen’s *d* >1.9) in all cases. Additionally, we monitored destruction of targets in co-culture and found that our survival-optimized system allowed for complete preservation after 24-hour exposure to cytotoxic effectors. With this bioplatform, we anticipate investigators will be empowered to rapidly express and characterize T cell receptor responses, generate new knowledge regarding the patterns of T cell receptor recognition, and optimize novel therapeutic T cell receptors for improved cytotoxic potential and reduced cross-reactivity to undesired antigenic targets.

## Background

T cells are a crucial component of the vertebrate immune system responsible for driving adaptive surveillance and removing external pathogens and mutated cells from the body. The T cell compartment mediates adaptive immunity against potential threats by producing a repertoire of millions of unique clones, each armed with a distinct surface-expressed T cell receptor (TCR) generated via programmed somatic rearrangement of a set of germline-encoded TCR genes. Each of these semi-randomized TCR proteins, in turn, is capable of interacting with its own range of potential antigenic ligands. These ligands consist of short peptide sequences (epitopes), which are derived from the intracellular turnover of cellular proteins, processed, and displayed on the cell surface by major histocompatibility complex (MHC) proteins as peptide/MHC (pMHC) complexes. T cells mount immune responses when TCRs recognize pMHC complexes containing peptides derived from invading microorganisms, or altered peptides derived from mutated versions of normal proteins. Positive T cell recognition initiates numerous effector mechanisms aimed at directly destroying infected/mutated cells directly and/or recruiting other immune cells to the site.

T cells can control proliferation of pre-malignant tumors, however, this capacity is finite and, when overwhelmed, cancer can progress (1). Hence, therapeutic strategies have been under development for decades to artificially boost anti-cancer immunity, most often by promoting the function of T cells expressing tumor specific TCRs (2). Emerging modalities of immunotherapy have been founded on the principle of identifying and validating specific TCR sequences with known reactivity to target tumor-associated pMHC complexes, and harnessing their function in therapeutic classes such as adoptive TCR-T cell therapy (3) and soluble TCR- based T cell engagers (4).

TCRs with therapeutic potential can be directly isolated from patient-derived tissue by selective priming and expansion of tumor-responsive T cell clones *ex vivo* using therapeutic target peptides (5,6), or by sorting T cells with antigen-specific TCRs on the basis of binding to pMHC multimer reagents (7,8). Bulk or single cell TCR-seq methods can then identify T cell clonotypes from which candidate TCR therapeutics may be selected for further development. However, TCR-seq provides no information regarding the functional reactivity of any TCR sequences identified. While TCRs are understood to be precise and selective molecular sensors, they are also known to be highly promiscuous and able to respond to a wide landscape of potential peptide epitopes (9,10). In large part, this paradoxical interplay between selectivity and promiscuity is mediated by the biophysically complex mode of activation characteristic of the TCR: various phenomena such as serial triggering, kinetic proofreading, and allostery in the immune synapse all contribute to the ultimate response of a given T cell clone (11,12). Currently, *in silico* prediction of these properties is not adequate for assessment of TCR proteins with respect to therapeutic potential (13). Functional assessment of candidate therapeutic TCRs, with a broader view of the epitope space they are able to respond to, is necessary to characterize potency and cross-reactivity.

We previously described a novel *in vitro* function-based method to assay cytotoxic T cell (CTL) responses and extended the assay format to be compatible with high-throughput screening (HTS) of T cell receptors-of-interest (TOI) against large sets of synthetic DNA-encoded peptide epitopes (14) in a method we term T cell epitope sequencing (Tope-seq). Briefly, the approach is centered around a granzyme-B (GZMB)-sensitive Förster Resonance Energy Transfer (FRET)-based reporter transgene, which produces GZMB-cleavable ECFP-EYFP fusion protein that undergoes a shift in fluorescence properties (FRET-shift) upon cleavage by GZMB delivered to target cells by antigen-responsive CTLs in co-culture. FRET-shift is monitored in flow cytometry to detect cells actively targeted by the CTL population. The early read-out of cytotoxicity and target-centric encoding of the GZMB reporter are features which enable FRET-shifted target cells to be isolated by fluorescence activated cell sorting (FACS) and subjected to next-generation sequencing techniques to reveal the identities of minigene sequences eliciting reaction from the TOI.

The study of T cell populations from genetically diverse human donors requires advanced strategies for ensuring compatibility between antigen presenting cell (APC) MHC genotype and T cell MHC restriction, in contrast to studies in murine systems, for example, where congenic strains and TCR-transgenic organisms simplify tissue sourcing for functional assays. Further, it is not always possible to achieve the level of T cell expansion or enrichment necessary for HTS experiments such as Tope-seq. For example, chronically activated T cells or T cells subjected to the highly immunosuppressive tumor microenvironment are often permanently attenuated and unable to proliferate or function *ex vivo* (15). In other scenarios, TCRs-of-interest can be identified as potentially relevant to health and disease based on retrospective TCR-seq datasets (16) but cannot be directly used for *in vitro* testing due to unavailability of source material.

In this study, we describe a novel synthetic platform which circumvents common challenges associated with generating suitable human tissues for functional TCR assessment while demonstrating implementation of the GZMB FRET-shift based read-out of TCR response in a fully reconstituted setting. To achieve this, we leverage YT-Indy, a natural killer acute lymphoblastic leukemia cell line which has retained functional granzyme B/perforin (PRF) delivery (17), and K562, a chronic myelogenous leukemia cell line negative for class-I HLA expression but positive for β_2_ microglobulin expression and the class-I antigen processing and presentation pathway (18). We demonstrate that, when exogenously provided with a cognate set of recombinant TCR, HLA, and peptide-coding sequences, *de novo* antigen-specific function can be induced and detected upon co-culture. We then characterize the resulting GZMB/PRF induced apoptosis kinetics of the system and further engineer the cellular platform to delay target cell death due to cytotoxic effector cell responses, which enables recovery and characterization of granzyme-loaded target cells.

## Results

### CD3z is sufficient for antigen receptor signaling in YT-Indy cells

Previous literature indicates that YT-Indy cells, while being a potent cytotoxic effector towards some cancer cell lines, do not result in cytotoxic killing of K562 cells *in vitro* (17). Thus, we speculated that it could be possible to redirect the existing cytotoxic potential of the YT-Indy cell line towards K562 in an antigen-specific manner if a cognate immune receptor/antigen combination was recombinantly expressed in the cell pair. Moreover, we hypothesized that if the cytotoxic effects mediated by YT-Indy were predominantly mediated by granzyme B/perforin delivery to target cells, we would be able to detect antigen-specific response using our previously described GZMB activated FRET-shift flow cytometry assay.

As a first test, we produced a lentiviral vector encoding the GZMB-cleavable ECFP-EYFP fusion protein described in our previous work and used it to transduce unmodified K562 cells. We also transduced 721.221 cells, which are a gamma-irradiated variant of the B-lymphoblastoid cell line, 721, and is one of the cell lines previously documented to be susceptible to YT-Indy mediated cytolysis (19). The resultant K562.FRET and 721.221.FRET cell lines were FACS-purified to remove untransduced or non-productive transductants and co-cultured with YT-Indy cells prior to flow cytometry analysis. More than 50% of target cells were FRET-shifted in YT-Indy + 721.221.FRET co-cultures after just 1 hour while only background level shifting was visible in YT-Indy + K562.FRET co-cultures even after 12 hours (Fig. 1a), indicative of GZMB delivery from YT-Indy to 721.221 cells but not to K562 cells. We also observed cell counts in flow cytometry of post co-culture target cells relative to time-zero (T_0_) co-culture controls for each experiment, assembled by preparing a parallel unmixed set of effector and target cells to be combined at the end of co-culture and immediately prior to flow cytometry sample preparation. Consistent relative count measurements are achievable in flow analysis by holding input cell numbers, resuspension volumes, flow rate, and acquisition times fixed across all samples (Supplementary Fig. 1). By comparing relative viable target cell counts obtained in experimental and matched T_0_ controls, we determined that no cell dropout due to lysis occurred in K562 co-cultures (Fig. 1b). From these data, we conclude that the K562 cells do not natively trigger a GZMB/PRF response in YT-Indy and that no alternative mode of cellular cytotoxicity occurs in this cell pair since all input target cells remain viable after exposure to effectors.

**Figure 1.**
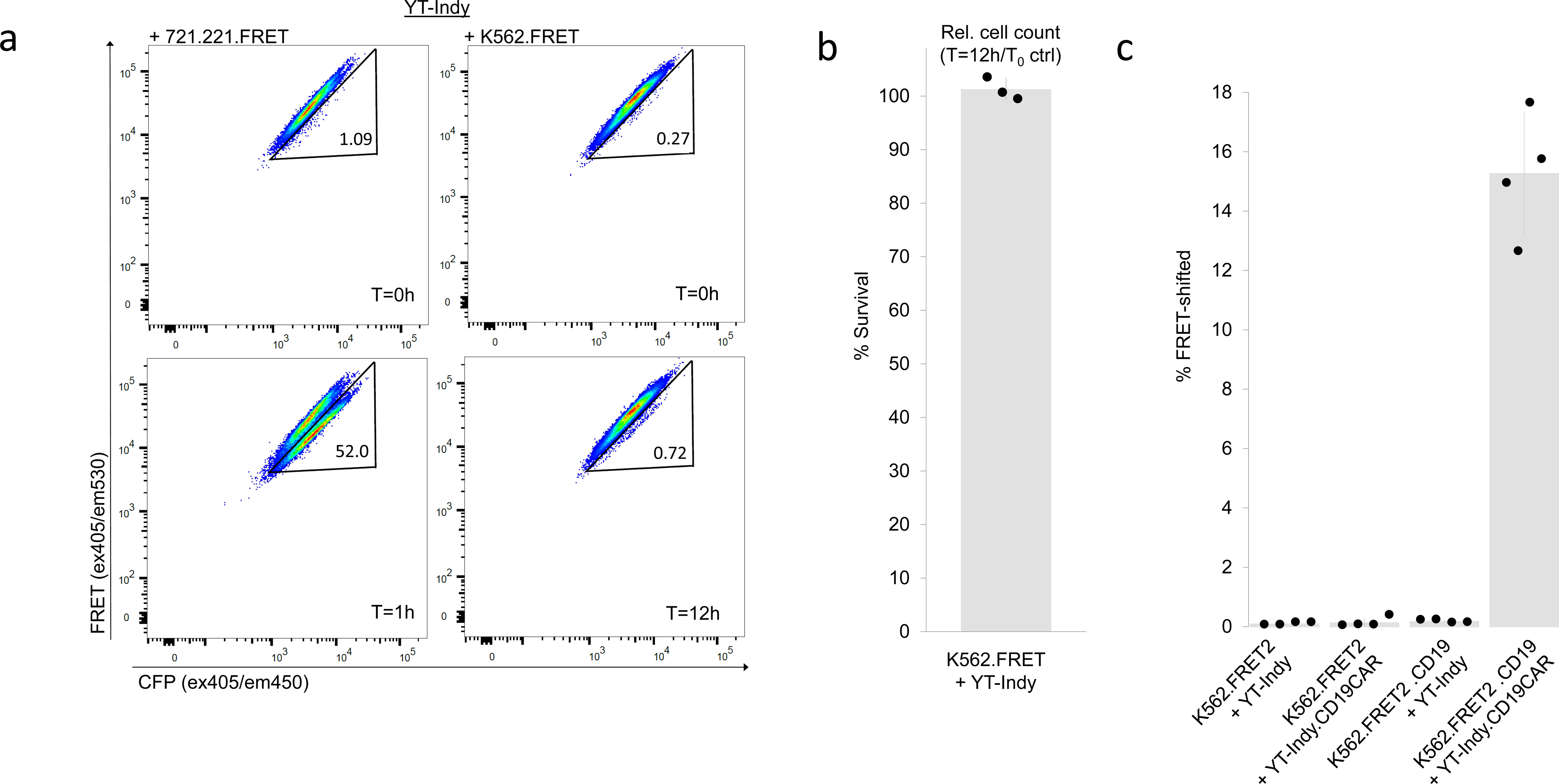
YT-Indy cells are GZMB/PRF competent but naturally spare K562 cells. (a) YT-Indy cells were co- incubated with either 721.221 cells or K562 cells, each of which was modified by lentiviral transduction to express GZMB-cleavable ECFP-EYFP FRET-reporter. Co-cultures were assembled at 1:1 effector:target ratios and incubated for the time intervals specified. Parallel T=0h (T_0_) controls were prepared for each co-culture condition to assess baseline FRET-shift signal and measure cell loss in co-cultures due to effector cell activity. (b) The ratio of cell counts from live, single-cell targets gate in flow cytometry (after holding input cell number, resuspension volume, flow rate, and acquisition time fixed for all samples) between K562.FRET/YT-Indy 12 hour co-cultures and T_0_ controls is shown (n=3, bar height and error bar represent mean ± standard deviation). (c) K562 cells modified to express FRET2 reporter (GZMB-cleavable CyPet-YPet fusion protein) were either additionally modified with CD19 protein coding sequence or left as CD19^-^. YT-Indy cells or YT-Indy modified with an FMC63-41BB-CD3ζ anti-CD19 CAR were each co-incubated with both prepared K562.FRET2 and K562.FRET2.CD19 target lines at 1:1 effector:target ratios for 4 hours. Percent FRET-shifted values from four independent replicate experiments are plotted (bar height and error bar represent mean ± standard deviation).

After confirming the feasibility of using our GZMB FRET-reporter as a sensor of YT-Indy cytotoxic response, we next addressed whether the cell line sufficiently expressed proximal signaling proteins necessary to result in GZMB/PRF response upon introduction and activation of exogenous antigen receptors to known cognate antigen. To this end, we used chimeric antigen receptors (CAR) as a preliminary test case. Chimeric antigen receptor constructs are composed of affinity domains consisting of single-chain antibody variable (scFv) fragments fused to intracellular T-cell signaling domains, the most critical one for cytotoxic response being CD3ζ (20). The configuration of the anti-CD19 BB ζ CAR we use here is based on a previously described design (21). The CAR transgene was lentivirally delivered to YT-Indy cells. In parallel, CD19 coding sequence was transduced into K562 expressing an alternate version of the GZMB FRET-reporter (termed FRET2) consisting of CyPet and YPet (22) fluorescent domains in place of the ECFP and EYFP proteins (respectively) that we previously reported (14).

Surface expression of CD19 in transduced K562 cells and anti-CD19-CAR in YT-Indy cells was confirmed by mAb staining and used to sort transduced cells to purity. The resultant cell lines were co-cultured and analyzed by FRET- shift flow cytometry (Figure 1c). The results show that YT-Indy.CD19-CAR cells initiate a response only when both the CD19 protein and the CD19-CAR are present in their respective target/effector cells in co-culture, indicating that the minimal signaling domains present in the intracellular region of the CAR design are sufficient to integrate with the downstream signaling pathways used by YT-Indy cells to mediate GZMB/PRF delivery to K562. Since the particular CAR design used in this experiment also encoded co-stimulatory 4-1BB intracellular domains alongside the canonical activation domain, CD3ζ, we also created a sequence-engineered version of the CD19-CAR transgene with an inserted premature stop codon downstream of the 4-1BB domain and upstream of CD3ζ, resulting in a truncated CD19-CAR lacking a CD3 signaling motif. When co-cultured with CD19-expressing K562 cells, we were able to confirm the intracellular 4-1BB domain does not appreciably contribute to YT-Indy degranulation and that CD3ζ is indispensable for driving YT-Indy.CD19-CAR based GZMB/PRF function, (Suppl. Fig. 2).

### TCR complexes can be reconstituted in YT-Indy cells

We next sought to reconstitute antigen-specific TCR function in the cytotoxic YT-Indy chassis. To test this, we used a previously described TCR sequence discovered in the ascites of a high-grade serous ovarian carcinoma patient (23). This TCR was found to be specifically responsive to a mutational neo-epitope arising from an L25V substitution in the amino acid sequence of hydroxysteroid dehydrogenase–like protein 1 (HSDL1). We initially attempted to generate HSDL1 mutant-reactive reconstituted CTL (rCTL) under the hypothesis that providing the TCRαβ chains of this clonotype, along with CD8α and CD3ζ domains would be minimally sufficient for redirecting YT-Indy cytotoxicity towards HSDL1 mutant epitope-bearing K562. Previous reports have shown that CD8α are able to form homodimers capable of binding MHC-I (24) while the activity of the YT-Indy.CD19-CAR suggested that CD3ζ domains are sufficient to trigger effector degranulation. To test this, a TCRαβ-RFP transgene and a CD8α/CD3ζ transgene were delivered to YT-Indy cells using lentivectors. Double-transduced cells robustly expressed surface CD8α and intracellular RFP (indicative of successful TCR transduction and transcription), but no surface TCRαβ was detected (Fig. 2a).

**Figure 2.**
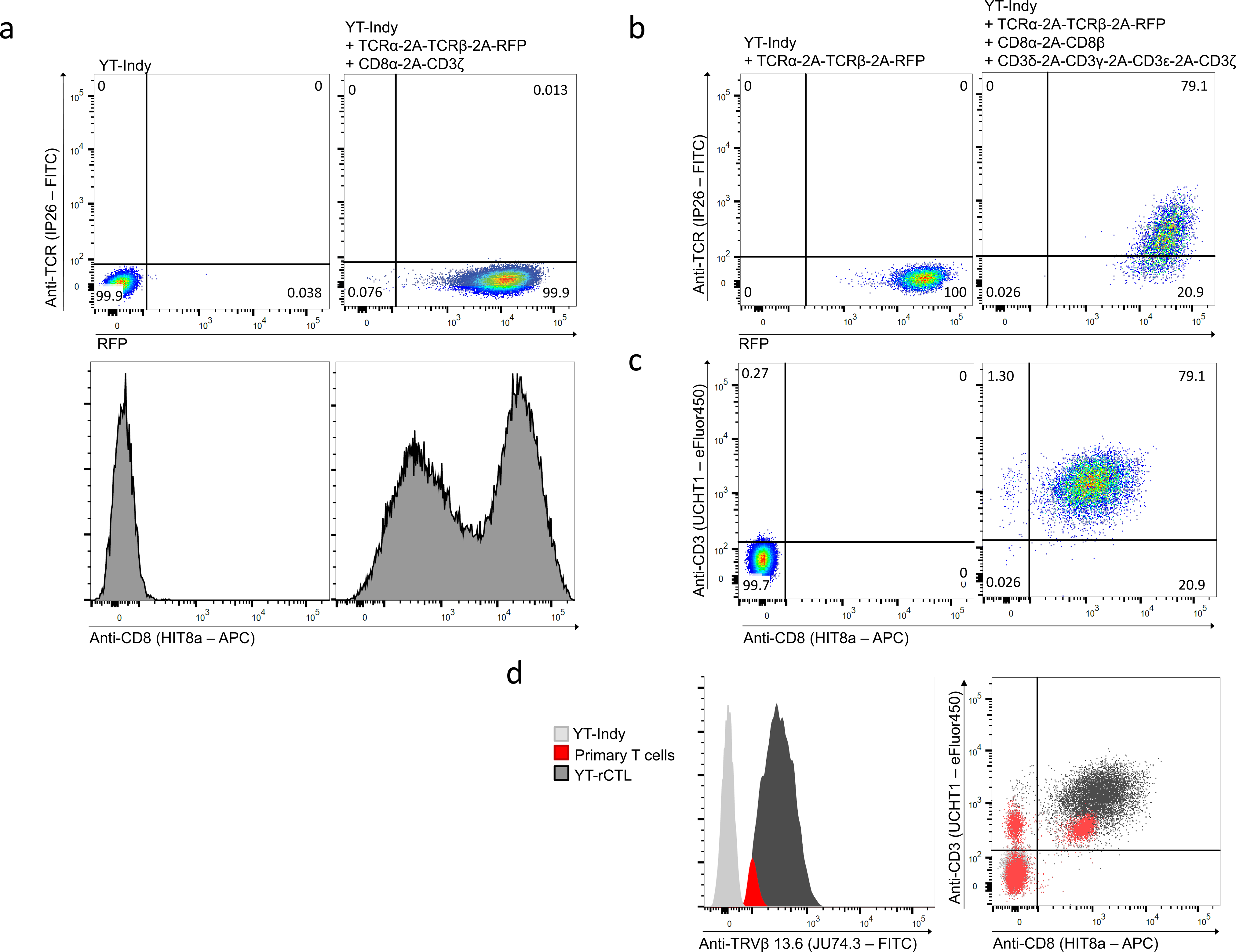
Recombinant expression and surface detection of T cell receptors in YT-Indy cell line. (a) Unmodified YT-Indy or YT-Indy cells transduced with lentivirus encoding HSDL1 L25V mutant-reactive TCRαβ followed by bicistronic CD3ζ/CD8α lentiviral helper cassette were analyzed by flow cytometry. Analysis confirms the robust expression of an RFP marker included with the TCRαβ transgene but no detection of TCR at the cell surface by antibody surface staining with FITC-conjugated anti-human TCR IP26 mAb clone (Abcam). Expression of CD8α at the cell surface was confirmed by staining with APC-conjugated anti-human CD8 HIT8a mAb (Biolegend). (b) YT-Indy cells previously transduced with mutant HSDL1-reactive TCRαβ encoding construct were double-infected with two helper cassettes designed to deliver complete CD3 and CD8 complexes. Detection of FITC fluorescence, indicative of surface TCRαβ binding, was observed to co-occur with robust RFP expression. In contrast, no TCR protein was detectable in YT-cells lacking helper cassettes despite expressing the RFP marker of transgene integration. (c-d) The TCR^+^ CD8^+^ double-positive population was isolated by FACS from triple transduced YT-rCTL shown in (a) and further characterized by surface expression. Strong expression, on the basis of mean fluorescence intensity (MFI) relative to primary human T cells comparator, was detected in CD8α, CD3ε, and TRBV channels. Surface staining of CD3ε was conducted using eFluor450-conjugated mAb clone UCHT1 (eBioscience). Surface staining of TCRβ was conducted using FITC-conjugated mAb clone JU74.3, specific for the known Vβ segment used by the HSDL1 TCR (Beckman Coulter).

Original characterization of the YT-Indy cell line showed that they are CD3^-^ by surface antibody staining but retain germline configuration of their native TCR loci (17). Since all CD3 subunits are required for stable assembly and surface trafficking of the TCR complex (25–27), it was unclear whether the CD3^-^ phenotype was due to lack of TCR expression and/or lack of expression of one or more CD3 subunits. Our initial result showed that exogenous rearranged TCR was inadequate for assembly of surface detectable TCR, therefore, we inferred YT-Indy cells lack native expression of CD3δ, ɣ, and ε. We confirmed this by performing RNA-seq and comparing relative transcript levels of CD3 subunit genes between YT-Indy cells and primary T cells (Supplementary data). This data revealed >200-fold higher normalized count of CD3δ and CD3ɣ and >2-fold higher normalized count of CD3ε and CD3ζ in primary T cell than YT-Indy.

Thus, we revised our approach to reconstituting TCR in YT-Indy cells by generating two additional lentivectors, one encoding all four CD3 subunits separated by distinct 2A signal sequences, and the other encoding the CD8α and CD8β genes separated by a 2A sequence. Upon co-transduction of the YT-Indy.HSDL1(L25V)-TCR intermediate cell line from the previous section, surface expression of the TCR complex was detected by antibody staining (Figure 2b,c). This newly generated YT-Indy reconstituted cytotoxic cell line (YT-rCTL.HSDL1) was observed to have CD8, CD3, and TCR surface levels higher than those observed on primary human CD8^+^ T cells (Figure 2d).

An unexpected observation during this process was the remarkable amenability of the YT-Indy cell line for lentiviral transduction, which facilitated the efficient creation of the variant cell line modified with three separate viral vectors. Comparison of titering data obtained from measuring the same batch of TCRαβ-2A-RFP viral preparation in either YT-Indy cells or primary T cells undergoing active expansion revealed that approximately 1x10^3^-fold fewer infectious units of lentivirus were required to productively infect the YT-Indy population as were needed for primary T cell transduction (Suppl. Fig 3). Therefore, in addition to serving as a cell chassis into which individual TOI may introduced for recombinant expression, we suggest that the heightened transduction efficiency in YT-Indy cells may also be an enabling property for assembling large libraries of recombinant TCR from relatively small preparations of lentivirus.

### YT-rCTL produce antigen-specific granzyme/perforin responses

Upon confirming that TCR expression could be achieved in the YT-Indy base cells, we set out to determine if GZMB/PRF cytotoxic function could then be redirected towards cognate peptide-MHC ligands displayed on K562 cells. We transduced K562 with HLA-C*14:03, which restricts the HSDL1 L25V reactive TCR, to generate a new line, K562.C1403. Surface expression of MHC protein on purity-sorted K562.C1403 was equivalent to surface MHC levels observed on primary PBMC controls from healthy donor samples (Suppl. Fig. 4). K562.C1403 was then transduced with cassettes comprising a wild-type (wt) or L25V-mutated minigene followed by a 2A site and the GZMB-cleavable FRET-reporter to produce the KFRET.C1403.HSDL1^4-43^.wt and KFRET.C1403.HSDL1^4-43^.L25V lines, respectively. Each of these newly prepared sAPC lines were co-cultured with YT-rCTL.HSDL1 effectors from the previous section for 12 hours and then analyzed by flow cytometry using the gating template illustrated in Suppl. Fig. 5. Robust FRET-shift signal was detected in L25V minigene-expressing cells when co-cultured with YT- rCTL.HSDL1 cells, while no appreciable signal was detected in targets expressing wt minigene (Figure 3a,b).

**Figure 3.**
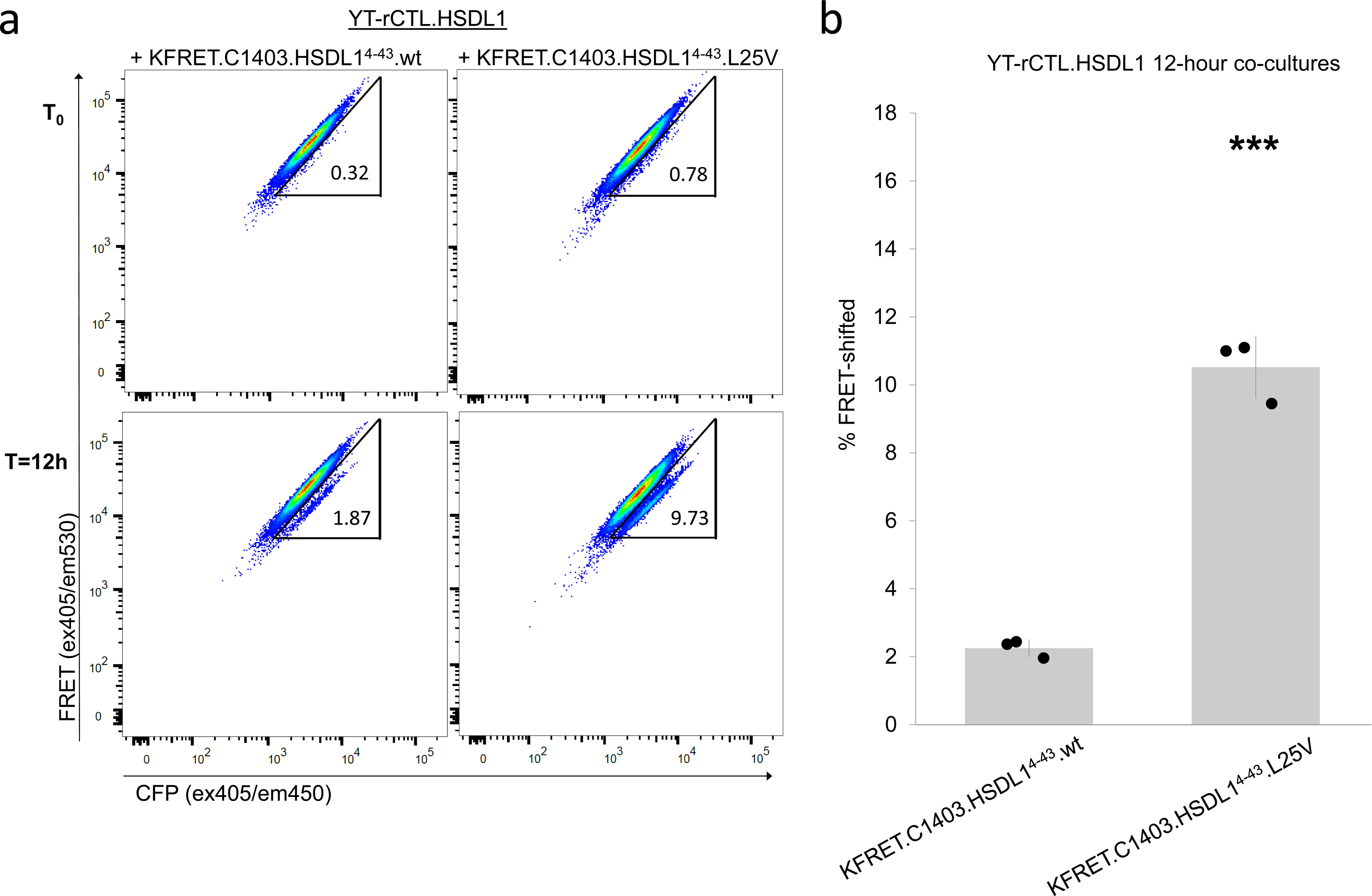
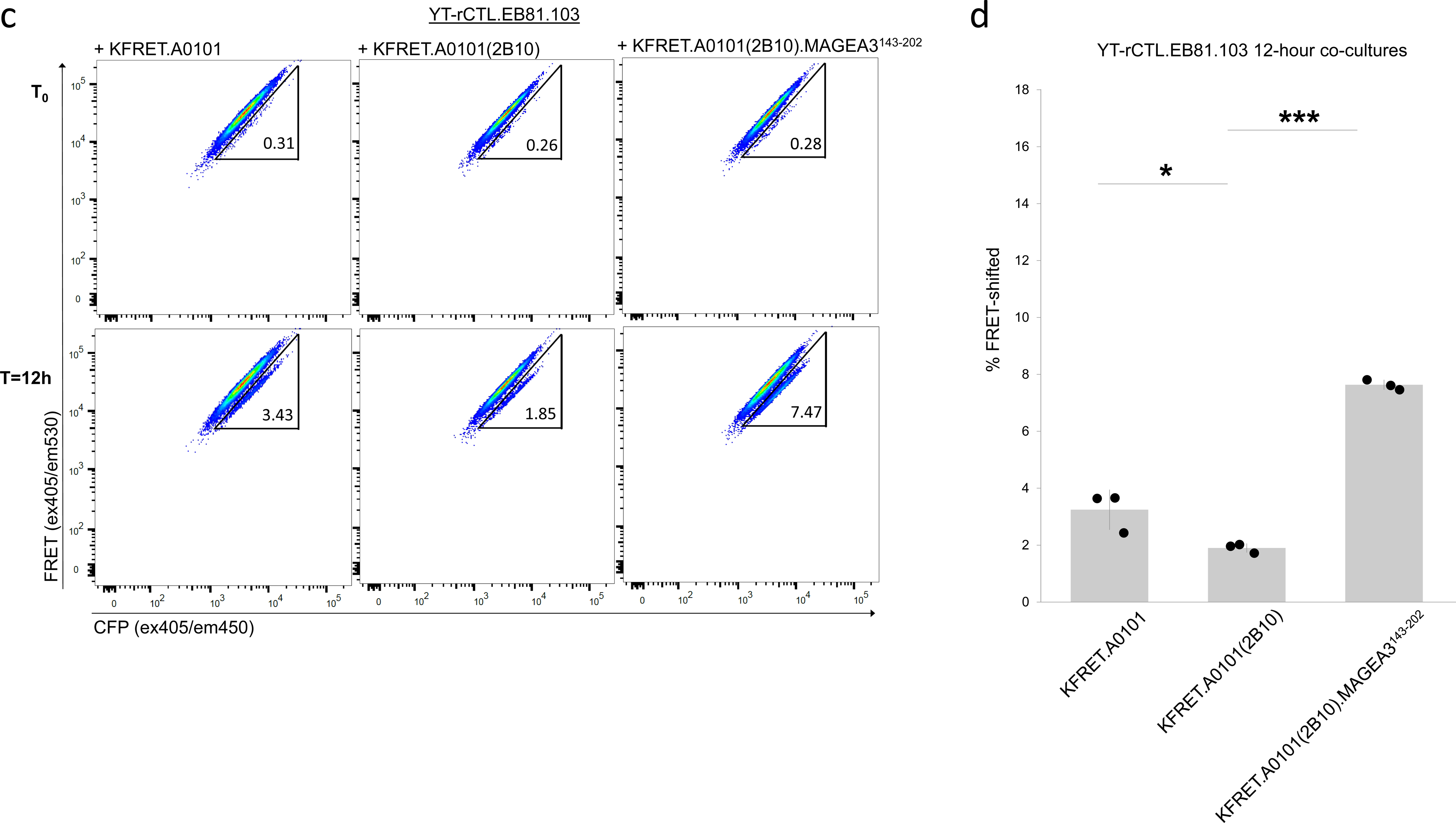
Antigen-specific induction of GZMB delivery from YT-rCTL to KFRET sAPC. (a-b) YT-rCTL expressing the HSDL1 L25V mutant-reactive TCR were co-cultured with mutant minigene-expressing KFRET.C1403.HSDL1^4-43^.L25V cells or the wild-type counterpart KFRET.C1403.HSDL1^4-43^.wt cells at 1:1 effector:target ratios for 12 hours. Matched T_0_ control co-cultures were also assembled and analyzed for each condition. Representative assay responses are shown in (a). Data from three independent replicate experiments indicate statistically significantly higher percent FRET-shift signal in mutant minigene co-cultures compared to wild-type minigene (*** p < 0.005 by unpaired two-sample t-test) (b). (c-d) YT-rCTL expressing the MAGEA3 reactive EB81.103 TCR were co-cultured with KFRET.A0101 cells, KFRET.A0101(2B10) epitope knockout cells, or epitope re-introduced KFRET.A0101(2B10).MAGEA3^143-202^ cells at 1:1 effector:target ratios for 12 hours. Matched T_0_ control co-cultures were also assembled and analyzed for each condition. Representative assay responses are shown in (c). Data from three independent replicate experiments indicate statistically significantly higher percent FRET-shift signal in KFRET.A0101 (* p < 0.05 by unpaired two-sample t-test) and KFRET.A0101(2B10).MAGEA3^143-202^ (*** p < 0.005 by unpaired two-sample t-test) compared to MAGEA3 epitope knockout cells All bar heights and error bars represent mean ± standard deviation.

We further validated antigen-specificity in the YT-rCTL/K562-FRET system using a melanoma-reactive TCR, clone 103 from patient EB81, here shortened to EB81.103 (28), responsive to a previously characterized peptide epitope (EVDPIGHLY) derived from the human MAGEA3 protein presented in the context of the HLA-A*01:01 allele. We created a new version of YT-rCTL cells by transducing the EB81.103 TCR into pre-made CD8ɑβ^+^, CD3δɣεζ^+^ TCRɑβ^-^ YT-Indy cells (to create YT-rCTL.EB81.103) and a new K562 subline encoding the HLA-A*01:01 sequence bicistronically linked to the GZMB FRET-reporter (to create KFRET.A0101) (Suppl. Fig. 4). As the K562 cell line expresses the MAGEA3 oncogene naturally (29), we disrupted the EVDPIGHLY epitope within the endogenous MAGEA3 gene using CRISPR/Cas9 editing (Suppl. Fig. 6a). Single cell cloned MAGEA3 mutant K562.A0101 (clone 2B10), when co-cultured with YT-rCTL.EB81.103 showed a significant reduction in FRET- shift signal consistent with ablation of the endogenous epitope. Reintroduction of agonistic epitope by lentiviral transduction of a minigene encoding the MAGEA3^143-202^ peptide fragment into 2B10 knockout cells resulted in rescue and significant increase of FRET-shift signal relative to endogenous antigen (Figure 3c,d). Notably, FRET- shift signal developed in YT-rCTL co-culture with KFRET.A0101(2B10).MAGEA3^143-202^ closely mirrored that from primary TCR-T cell effectors (Figure 3c,d/Suppl. Fig. 6b).

Taken together, these data indicate that reconstitution of the TCR/CD3 complex in the GZMB/PRF competent natural killer cell-origin immortalized cell line, YT-Indy, induces antigen-specific cytotoxic responses against K562 based target cells only in the presence of an MHC and peptide epitope combination cognate to the recombinant TOI. We show in the contexts of mutational neoantigens or overexpressed self-antigens, and in different HLA alleles, that the YT-rCTL/K562-FRET system is a highly selective platform for biologically-relevant detection and measurement of TCR interactions.

### YT-rCTL are more potent than primary TCR-T cells

We previously established that there is a delay between GZMB delivery to targets and their subsequent destruction due to programmed cell death. This allows for a “safe-sorting” window for GZMB FRET-shift assay representing the period of time in which cells attacked by cytotoxic T cells can be isolated by FACS. To characterize co-culture kinetics specific to the YT-rCTL/KFRET system, we selected an alternative, affinity-enhanced TCR for recombinant expression and testing. This TCR, called a3a, is based on the EB81.103 clonotype but has been engineered to contain variants that increase affinity for the MAGEA3-(EVDPIGHLY)/HLA-A*01:01 pMHC complex. The original description of the a3a TCR (30), reported considerably stronger pHLA binding than the relatively weak wild-type EB81.103 counterpart and stronger cytotoxicity towards tumor targets *in vitro*. Clinical development of the a3a TCR was halted due to the death of two patients from cardiotoxicity, caused by cross-reactivity of a3a to an epitope in the titin protein (TTN) expressed in cardiomyocytes. (31). We chose to study target cell dropout kinetics in the context of the a3a TCR in order to characterize the most extreme condition likely to be encountered in real-world application of the YT-rCTL/KFRET system.

We transduced CD8ɑβ^+^, CD3δɣεζ^+^ TCRɑβ^-^ YT-Indy cells with the a3a TCR and tested them in 12 hour co-cultures with KFRET.A0101(2B10) and KFRET.A0101(2B10).MAGEA3^143-202^ sAPCs. As expected, YT-rCTL.a3a effector cells resulted in a substantial increase in FRET-shift signal relative to their wild-type counterparts (Figure 3c,d/Suppl. Fig. 7). Incidentally, we also observed that YT-rCTL.a3a-TCR resulted in a reduced but still substantial FRET-shift signal against 2B10 MAGEA3 knockout cells compared to the parental line. This was unanticipated since K562 cells do not express the TTN protein previously elucidated as the source of cross-reactive epitope mediating clinical failure, indicating that other uncharacterized cross-reactive epitopes likely exist in the K562 base cell line. Interestingly, parallel experiments of primary PBMC-derived a3a TCR-T cells showed virtually no change in signal between the two sAPC lines (Suppl. Fig. 7), suggesting that YT-rCTL responses could be additive with increasing numbers of distinct cognate pMHC ligands present, whereas the primary TCR-T cells may hit a functional ceiling.

After initial testing we conducted time course experiments to monitor the evolution of the FRET-shift signal as well as to measure the survival time of KFRET.A0101.MAGEA3^164-179^ target cells in co-culture upon exposure to YT- rCTL.a3a cytotoxic effectors. Percent survival was measured by comparing the count of target cells remaining in co-culture samples at each time point relative to a matched T_0_ control sample. Our results indicated immediate and substantial loss of target KFRET.A0101.MAGEA3^164-179^ cells. Peak FRET-shift signal (>40%) was achieved 8-12 hours after initiation of co-culture, however, by this point >60% of input targets had been destroyed and dropped out of culture (Fig. 4a). This result was a departure from previous work we had conducted characterizing GZMB apoptosis kinetics of co-cultures where we had noted that, using primary cytotoxic T cells, FRET-shift signal accumulated in target cells until reaching a peak which preceded the onset of a sudden, sharp decline in target cell count due to late-stage apoptosis (14). Indeed, in the present study we observed <10% dropout of target cells after 12 hours when K562-based sAPC were co-cultured with primary a3a TCR-T cells, suggesting that YT-rCTL based killing dynamics deviate significantly from those of normal cytotoxic T cells (Fig. 4b).

**Figure 4.**
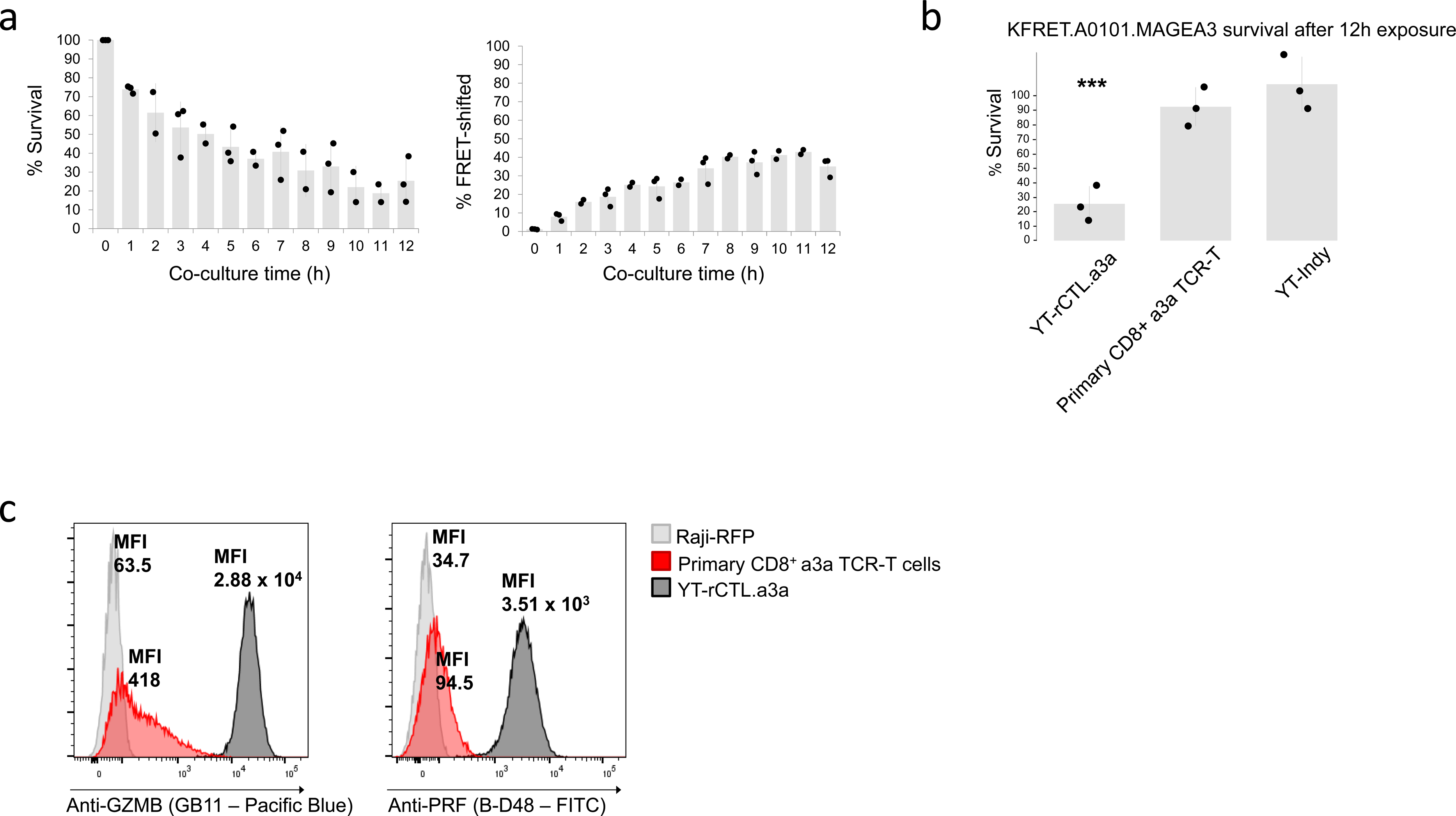
Monitoring target cell killing in FRET-shift flow cytometry of YT-rCTL/KFRET co-cultures. (a) Time course assessment of YT-rCTL.a3a + KFRET.A0101.MAGEA3^164-179^ co-cultures. Reactions were assembled by plating 2 x 10^5^ of each cell line into 2 separate wells of a U-bottom 96-well plate. An effector/target well pair was prepared for each of the 12 time points investigated and left unmixed until co-culture periods were initiated at the appropriate time prior to flow cytometry by combining effector and target wells and re-distributing co-cultures to both wells. Flow cytometry was conducted on all samples in a single batch at experiment conclusion. Percent FRET- shift signal and percent survival (live target cell counts normalized to matched T_0_ control) are displayed. Data are from three independent replicate experiments. (b) 12-hour YT-rCTL.a3a + KFRET.A0101.MAGEA3^164-179^ experiments were compared to 12-hour co-cultures of the same targets against either unmodified YT-Indy cells or primary cytotoxic T cells freshly isolated by FACS purification CD8^+^ CD4^-^ CD56^-^ T cells from healthy donor PBMC, activated by plate-bound anti-CD3/28 stimulation, and transduced with a3a TCR lentivirus. Unmodified YT-Indy cells were co-cultured with MAGEA3 targets at a 1:1 E:T ratio. Primary a3a TCR-T cells were co-cultured at a 1.5:1 ratio (to compensate for TCR transduction efficiency, which ranged from 57%-68% across replicates). Data are from three independent replicate experiments (primary TCR-T cells were freshly prepared for each experiment using different donors each time). YT-rCTL.a3a resulted in statistically significant (*** p < 0.005 by unpaired two-sample t-test) dropout of target cells after 12 hours when compared to both negative control conditions by two-sample unpaired t-test. (c) Primary CD8 a3a TCR-T cells, YT-rCTL.a3a, and Raji cell negative controls were fixed, permeabilized, stained with either anti-granzyme B or anti-perforin monoclonal antibody (Pacific Blue- conjugated clone GB11 and FITC-conjugated clone B-D48, respectively; Biolegend). The steady-state MFI of GZMB and PFR, as measured by intracellular staining and flow cytometry, are labeled.

### Genetic interventions as a strategy to protect K562 sAPC against destruction

Based on the above time course results, we concluded the YT-rCTL/KFRET platform would require further modification in order to capture cells containing potentially important epitopes for identification before they are lost from culture. We also noted in our preliminary experiments that the FRET-shifted fraction of K562 targets remaining viable after 12-hour co-culture, separated from effectors by FACS, were capable of recovering and proliferating in normal cell culture (Suppl. Fig. 8). Therefore, we speculate that increasing the proportion of targets surviving GZMB exposure during co-culture could also provide a means for preserving and propagating antigen-encoding targets indefinitely after co-culture, allowing for additional flexibility and capability in characterizing these populations. We pursued this on two fronts: attenuating the potency of the YT-Indy cell by knocking down expression of PRF and armoring K562 sAPC against GZMB-induced apoptosis by overexpression of anti-apoptotic genes.

Regarding attenuation of YT-Indy potency, characterization by intracellular cytokine staining flow cytometry of the base YT-Indy cell line revealed that at rest these cells store approximately 70-fold more GZMB and 40-fold more PRF than primary cytotoxic T cells (Fig. 4c). By RNA-seq, we detected 10-fold more *GZMB* transcript and 5-fold more *PRF1* transcript in YT-Indy cells relative to primary activated cytotoxic T cells (Supplementary Data). Our rationale for targeting only PRF for gene knockdown is based on previous work illustrating that supralytic concentrations of PRF result in rapid necrotic cell death independently of GZMB (32). We reasoned that shifting the balance of cell death progression away from necrosis and towards apoptosis would enable reliable cell recovery and *post hoc* analysis of targeted sAPCs within a known time period (33,34). We chose to pursue attenuation by lentivirally transducing YT-rCTL.a3a cells with miRNA-expressing cassettes containing either of three different designed sequences (using the BLOCK-iT™ RNAi Designer, Invitrogen). Each of the three PRF miRNA lines was then purity sorted by surface staining and detection of inert truncated CD34 marker included in the miRNA cassettes; for each transduced line, a high-, medium-, or low-level gate was selected to recover three different expression groups per miRNA construct (Suppl. Fig. 9a). All prepared cell lines were tested in 4-hour co-cultures with KFRET.A0101.MAGEA3^160-183^ cells (Figure 5a). Our results indicated that there was no appreciable difference within miRNA groups between expression-level gated sub-populations. However, the PRF miRNA2 group resulted in a statistically significant increase in target cell survival without causing any decrease in concomitant FRET-shift signal. PRF miRNA 1 and 3 did not contribute to significant increase in survival.

**Figure 5.**
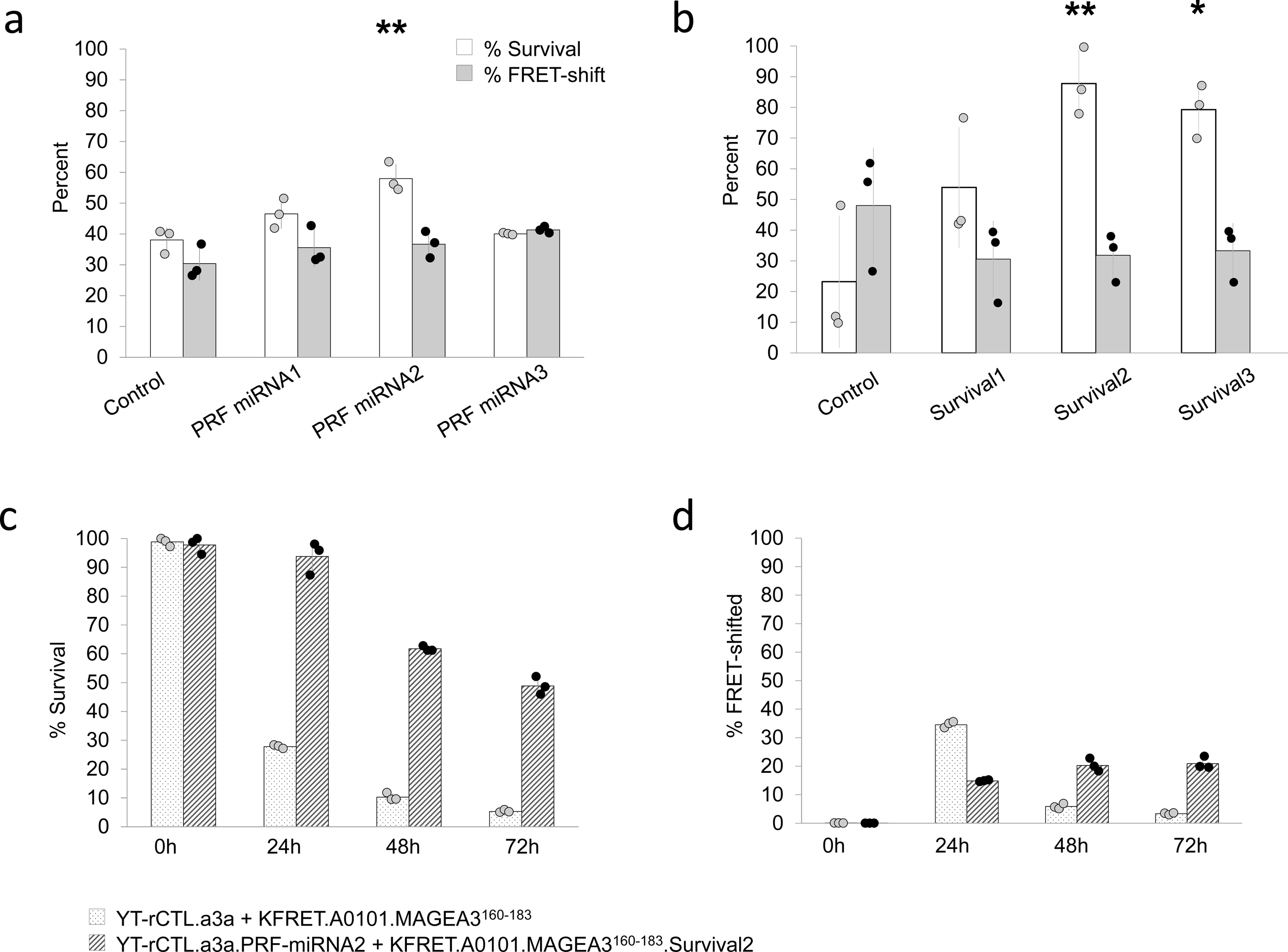
Genetic modification of YT-rCTL and/or K562 sAPC increases resistance to GZMB induced cell death. (a) Three separate YT-rCTL.a3a were produced, each expressing one of three different PRF targeting miRNAs. Each of these new YT-rCTL cell lines, along with a scrambled miRNA control, were co-cultured with KFRET.A0101.MAGEA3^160-183^ at a 1:1 E:T ratio for 6 hours. Percent FRET-shift signal and percent survival are displayed. Only miRNA2 resulted in statistically significant increase in percent survival relative to control (** p < 0.01 by unpaired two-sample t-test). FRET-shift signal was not statistically significantly different between control and any of the miRNA cassettes. (b) Three separate KFRET.A0101.MAGEA3^160-183^ cell lines were produced, each expressing one of the three different survival cassette configurations encoding the human genes XIAP, MCL1, and BCL2. Each of these new sAPC cell lines, along with an empty vector control, were co-cultured with YT-rCTL.a3a at a 1:1 E:T ratio for 18 hours. Percent FRET-shift signal and percent survival are displayed. Only Survival2 and Survival3 cassettes resulted in statistically significant increase in percent survival relative to control (** p < 0.01 and * p < 0.05 by unpaired two-sample t-test, respectively). FRET-shift signal was not statistically significantly different between control targets and any of the Survival cassette targets. (c, d) Four co-cultures of KFRET.A0101.MAGEA3^160-183^.Survival2 + YT-rCTL.a3a.PRF-miRNA2 or KFRET.A0101.MAGEA3^160-183^ + YT- rCTL.a3a were assembled and measured sequentially by flow cytometry at each of 0-, 24-, 48-, and 72-hour time points for percent survival (live target cell counts normalized to maximum count from each T_0_ control group) (c) and FRET-shift signal (d). No statistically significant change in percent survival was observed between T_0_ and 24-hour time point in the KFRET.A0101.MAGEA3^160-183^.Survival2 + YT-rCTL.a3a.PRF-miRNA2 condition. All data shown are results of three independent experiments.

To armor K562 sAPC, we selected three anti-apoptotic genes for use in constructing a survival cassette. Based on publicly available expression data, we noted K562 cells are devoid of natural *BCL2* expression (*BCL2* cell-line data available at v23.proteinatlas.org/ENSG00000171791-BCL2/cell+line (35)). Hence, we hypothesized that BCL2 protein replacement would result in decreased GZMB/PRF induced cell death. We also selected MCL1, another anti-apoptotic member of the BCL2 protein family, known to be directly degraded by GZMB (36). We reasoned that the addition of a constitutively active *MCL1* gene would function to replace protein turned over by GZMB activity. The third gene included in survival cassette designs was *XIAP*, a direct inhibitor of late-apoptosis executioner caspases (37). Three separate configurations for encoding all three selected proteins were designed. First, a polycistronic gene cassette consisting of all genes separated by 2A ribosomal-skipping sequences and linked to a truncated CD34 gene for use as a surface stainable marker of transduction. Since 2A sequences result in a C-terminal molecular scar in expressed protein (38) and since both BCL2 and MCL1 are membrane-anchored via C-terminal transmembrane domains, we suspected function of both proteins may have been impaired if placed upstream of a 2A sequence. Therefore, the second and third configurations were constructed using internal ribosomal entry site (IRES) sequences, instead of 2A sequences, from either encephalomyocarditis virus (ECMV) or the human *VCIP* gene (Table 1).

**Table 1.** Summary of survival cassette designs used in K562 armoring tests. Schematic description of gene-coding sequences inserted between lentiviral packaging elements of host transfer plasmid. All cassettes were placed under a constitutive MNDU3 promoter.

| Name | Cassette configuration |
| --- | --- |
| Survival1 | BCL2-P2A-XIAP-T2A-MCL1 |
| Survival2 | XIAP-P2A-BCL2-IRES(VCIP)-tCD34-T2A-MCL1 |
| Survival3 | XIAP-P2A-BCL2-IRES(ECMV)-tCD34-T2A-MCL1 |

Survival cassettes were transduced into KFRET.A0101.MAGEA3^160-183^ cells. Three purity-sorted cell lines, each expressing one of the survival cassettes, (Suppl. Fig. 9b) were then tested in 18-hour co-cultures with YT-rCTL.a3a cells (Figure 5b). The Survival2 and Survival3 cassettes led to statistically significant prevention of cell dropout but no statistically significant decrease in FRET-shift signal. Survival1 did not lead to significant protection of the target cells, suggesting that BCL2 is likely partially dampened by the presence of a 2A-derived C-terminal scar, as suspected, and is indispensable to Survival cassette function. Survival2 resulted in higher survival (>85% cells remaining) than Survival3 which, since *VCIP* IRES has been shown to result in higher expression of the downstream transcripts relative to ECMV IRES (39), is suggestive of a contributing role of the *MCL2* gene in the Survival cassette design.

We next investigated the combination of both strategies, using the best performing modification from each of the above-described arms. Co-cultures of YT-rCTL.a3a.PRF-miRNA2 and KFRET.A0101.MAGEA3^160-183^.Survival2 and were assembled and monitored with respect to percent survival and FRET-shift signal at 24-hour, 48-hour, and 72-hour time points. From these data, we determined that the survival cassette and PRF knockdown combination are indeed additive and nearly completely effective at averting cell death and dropout. On average, >93% of cells, relative to matched T_0_ controls, remained after 24-hour co-culture (Figure 5c). The percent FRET-shift signal was reduced, relative to unmodified controls, by a factor consistent with what had been observed in the initial survival cassette testing (Figure 5d). Strikingly, we also noted that despite nearly total destruction of target cells in the unmodified control conditions after 48 and 72 hours (<10% and <5% of cells remained, respectively), a significant fraction of target cells remained in the survival cassette/PRF knockdown condition after 48 and 72 hours (>60% and >45%, respectively).

Based on these data, we concluded that GZMB-loaded target cells could be preserved in culture within a safe-sorting window of up to 24 hours post co-culture initiation. This result expands the capability of the YT-rCTL/KFRET system beyond one-on-one assessment of TCR/pMHC pairs into functional screening of high diversity antigen libraries by FRET-shift based FACS isolation of targeted cells and characterization of these cells by DNA sequencing or other methods.

### Common markers of T cell activation in YT-rCTL

Having validated the YT-rCTL/KFRET system in discovering activating antigens encoded in sAPC, we explored its utility for TCR discovery by searching our RNA-seq dataset for T cell activation markers. Analysis was performed by comparing normalized read counts between samples of YT-rCTL.a3a or a3a TCR-T cells that were either co-cultured for 12 hours with KFRET.A0101.MAGEA3^160-183^ or not exposed to sAPC. Selected markers of activation are displayed in Table 2.

**Table 2.**
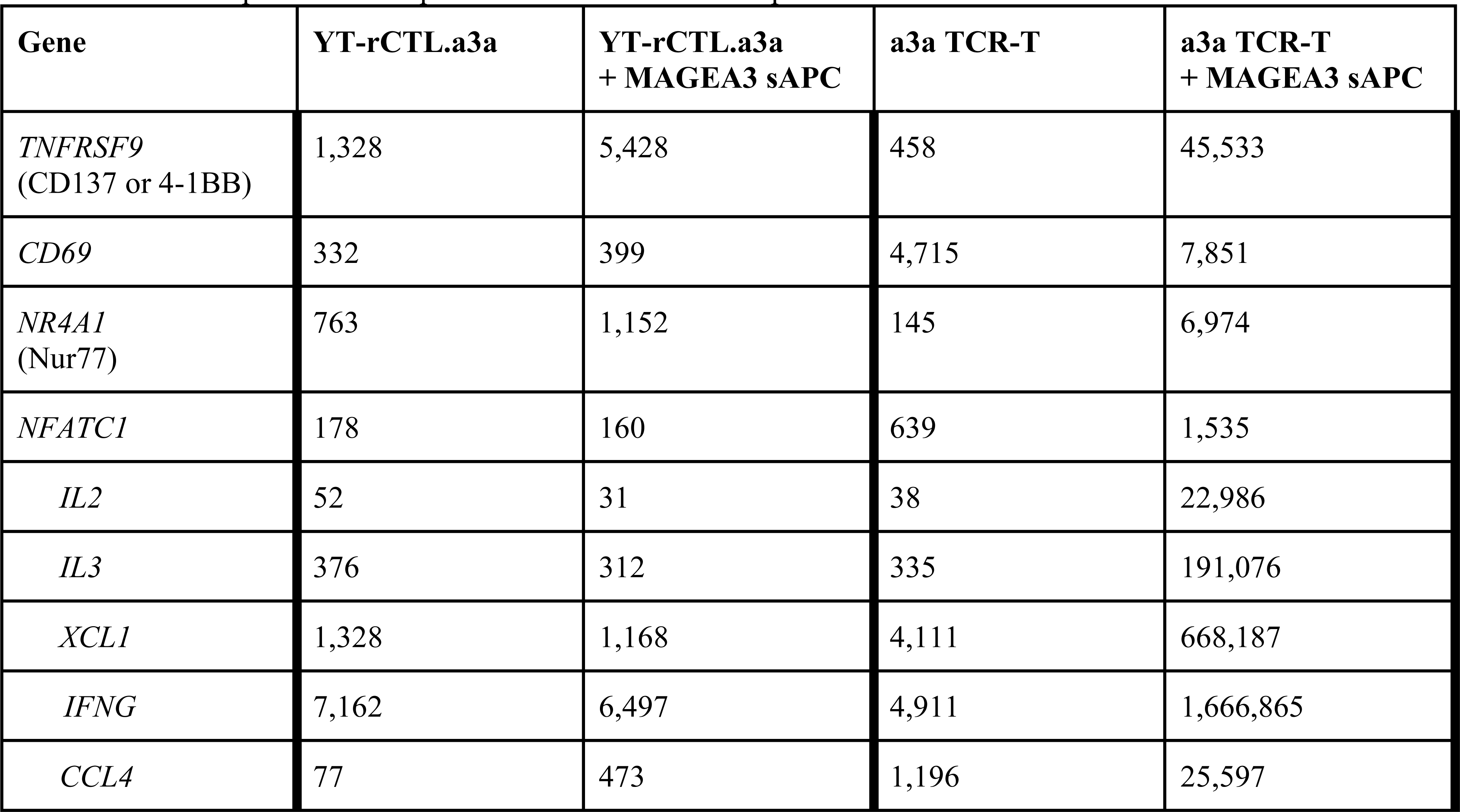
RNA-seq comparison of selected markers of T cell activation. Pairwise differential expression analyses between antigen-unexposed and post 12-hour co-culture samples of either YT-rCTL.a3a or a3a TCR-T cell format effectors. Values represent DESeq2 Normalized Counts. Group size for each condition was n=1.

The results indicated CD137 was upregulated 4-fold in YT-rCTL 12 hours after antigen exposure, but this was approximately 10-fold less than the degree of CD137 up-regulation seen in primary TCR-T. Direct assessment of CD137 by antibody staining on YT-rCTL and flow cytometric analysis showed that surface MFI could not be differentiated between T_0_ control and either 12-hour or 24-hour co-culture experiments (Suppl. Fig. 10), suggesting that this marker was not suitable for selection of responding YT-rCTL clones. Other conventional markers CD69 and NUR77 showed only modest upregulation in primary TCR-T cells, which would be expected since the 12 hour time point is beyond previously established peak signal for each these markers (40,41). In YT-rCTL.a3a these markers are expressed at relatively low levels and did not experience any appreciable upregulation in response to antigen, also precluding their use in this system.

Another widespread approach for monitoring specific activation in effector cells is the integration of reporter transgenes consisting of a detectable marker under the control of a T cell-specific transcription factor binding site (most commonly NFAT (42–45)). While differential expression in RNA-seq data cannot necessarily be taken as a measure of functional status of NFAT, since the NFAT protein can exist in inactive and active forms in the cell, we did note that NFAT expression was relatively low and unchanged upon antigen exposure in YT-rCTL cells. We further evaluated NFAT activity by monitoring known downstream genes under its control (46), which include IL2, IL3, XCL1, IFNG, and CCL4. In all of these downstream markers, large upregulation and high transcript counts were observable in primary TCR-T cells after antigen-exposure but these effects were not recapitulated in YT-rCTL effectors (Table 2), suggesting that NFAT-linked reporter systems may not be useful in YT-Indy cells.

Global gene expression in YT-rCTL and primary CD8^+^ T cells are shown in Figure 6. Positive markers of T cell activation are found to be highly expressed in the activated TCR-T cell population (Fig. 6a), whereas YT-rCTL cells do not appear to have a transcriptional signature of antigen response (Fig. 6b,c). We also assessed if it may be possible that genes with high expression in the absence of antigen could be downregulated during 12-hour co-culture, presenting marker loss as a possible readout of activation. However, no candidate markers were revealed by this analysis either (Fig. 6c). While unhelpful for TCR discovery, the aplasticity of the YT-Indy transcriptome may make these cells highly consistent and reproducible in co-culture and hence more amenable to antigen discovery and characterization approaches.

**Figure 6.**
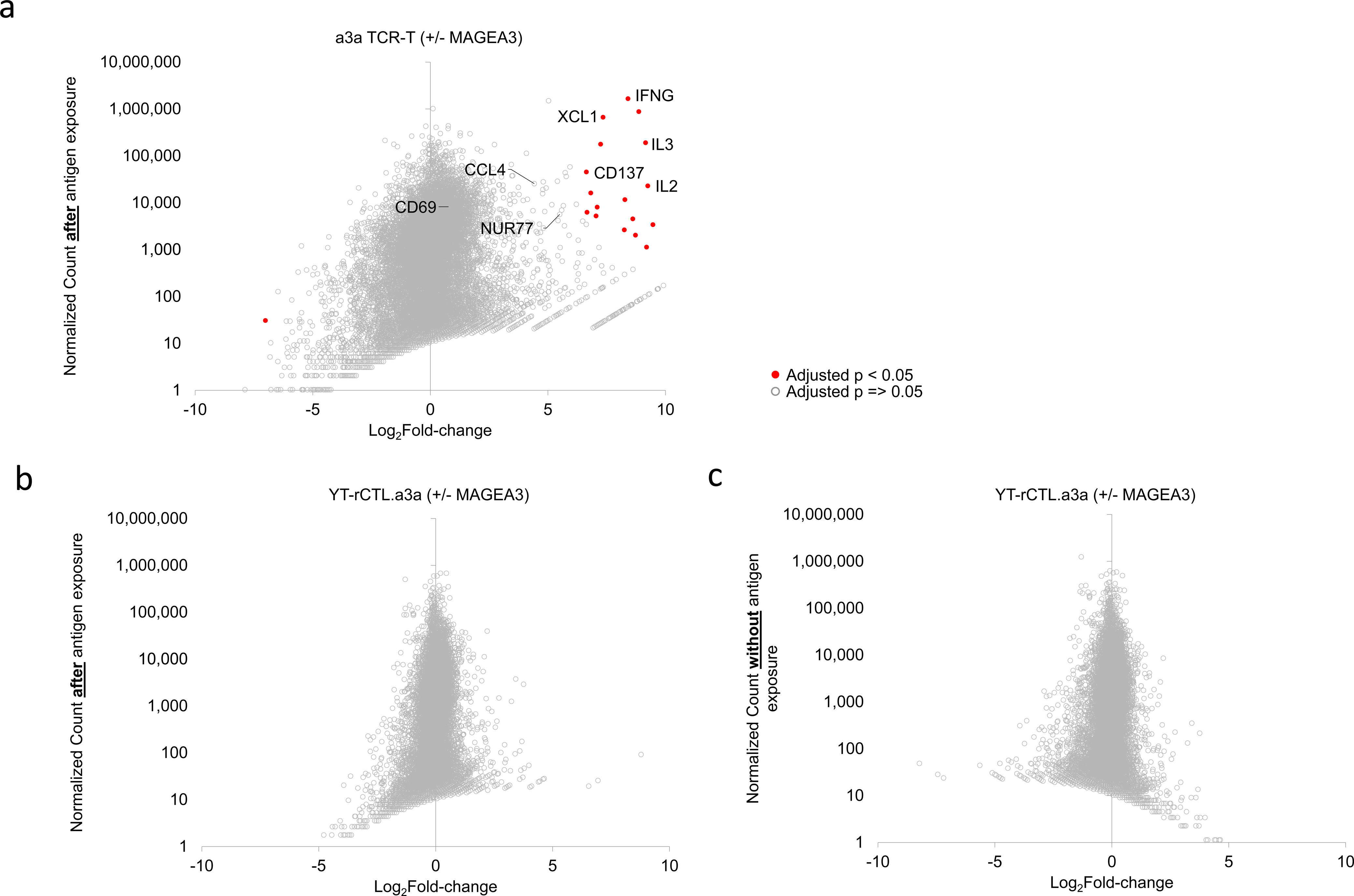
Transcriptional landscape change in TCR-T and YT-rCTL cells in response to antigen. Primary CD8^+^ TCR-T expressing a3a TCR (a) and YT-rCTL.a3a cells (b,c) were FACS sorted either after 12-hour co-culture with KFRET.A0101.MAGEA3^160-183^ sAPC or no co-culture. Flash-frozen cell pellets were subject to PE150 transcriptome sequencing on polyA-selected RNA. Pairwise comparisons between antigen-exposed and non- exposed conditions were conducted using DESeq2. Plots illustrate log_2_fold-change of all transcripts versus normalized read count of transcripts after exposure to sAPC (a,b) or normalized read count in the absence of sAPC (c). Genes referenced in Table 2 are labeled in (a). The standard deviation of log_2_fold-change in primary T cells = 1.61 and in YT-rCTL = 0.65; variance in log fold-change between both experiments was determined to be unequal by Levene’s test (p < 0.001). Eighteen data points were found to be statistically significantly differentially expressed in a3a TCR-T experiment (red filled circles denote p < 0.05 after Bonferroni correction), no statistically significant genes in YT-rCTL experiment detected.

Our RNA-seq survey of the transcriptional shift in YT-rCTL effectors is not a comprehensive view of all possibilities for developing an assay of effector response. Antigen-induced enzyme activity, post-translational modifications, or cellular re-localization of proteins cannot be discerned by RNA-seq yet could be leveraged into a novel reporter of activation specific for YT-rCTL. Further work will be needed on this front to address the opportunity to implement a novel effector-side readout to complement the target-centric GZMB FRET-shift assay.

## Discussion

Tools for functionally profiling T cell responses are crucial for decoding adaptive immunity and characterizing TCR therapeutics with respect to potency and cross-reactivity. The YT-rCTL/KFRET system we present here provides a stable and reproducible platform into which TCR-, HLA-, and peptide-coding DNA sequences of interest can be readily integrated and directly functionally tested. Central to this approach are two cell lines, CD8ɑβ^+^, CD3δɣεζ^+^ TCRɑβ^-^ YT-Indy cells and HLA^-^ K562 cells, which are non-reactive in co-culture until cognate TCR/pMHC are recombinantly added to the system.

In incumbent assays, effector cells are typically obtained by isolating, banking, and characterizing TOI in primary T cell clone format or recombinantly expressed TCR in primary cytotoxic T cell populations (*i.e.* in TCR-T cell format). In either of these cases, stimulated and expanded primary T cells are subject to senescence after a relatively short culture time, rapidly lose cytotoxic activity with serial re-stimulation and freeze/thaw cycles (47), are subject to substantial inter- or intra-donor batch-to-batch variability (48,49), and are only moderately amenable to genetic editing. Moreover, assays using TCR-T cell populations as effectors can be confounded by endogenously expressed TCR. Conversely, YT-rCTL can be kept in long-term cell culture, stably cryopreserved/thawed with no loss of performance, and are remarkably easily edited using lentiviral vectors. YT-rCTL lack endogenous rearranged TCR loci and, therefore, are also not subject to artifacts arising from endogenous TCR interactions. Effector cell chassis in the form of other immortalized cell lines could also be contemplated for use in the rCTL/sAPC platform described here, however, with limitations. The Jurkat cell line is the most well-established prior example of an alternative effector cell chassis in which TCR function can be characterized (50), however, it is CD4^+^ T cell in origin and does not naturally employ GZMB/PRF cytotoxicity. The popular TALL-104 and NK-92 cell lines, alternatively, are immortalized lines of T cell- and NK-origin, respectively, that do indeed retain intact GZMB/PRF functionality but both display broad TCR-independent cytotoxic tropism to a wide range of cells (51,52), limiting the choice of companion APC lines. Thus, despite the availability of other immortalized T and NK cell lines, YT-Indy is uniquely suited for rapid prototyping of TCR using the GZMB FRET-shift assay and the associated HTS configurations made accessible thereof.

The K562 cell line, besides being unsusceptible to intrinsic YT-Indy cytotoxicity, is advantageous as a synthetic APC chassis due to its natural MHC-null status, retained MHC class I antigen processing and presentation machinery, compatibility with contemporary genome editing techniques, and immortalization. Indeed, K562 has previously been established as a reliable artificial APC in related applications, such as *ex vivo* expansion of antigen- specific T cells (53–55). Other approaches to solving the requirement for HLA compatibility for functional TCR assays have also been described, each with significant limitations. In some cases, autologous cells from the same donor from which the TOI was derived is an attractive choice for generating a host cell population, however, these samples are rarely available, especially in the case of TCRs mined from *in silico* repertoire sequencing repositories. Allogeneic donor cells with a matched MHC allele-of-interest are another potential source of APC but with the limitation that the presence of foreign, non-TOI-restricted HLA alleles can introduce alloreactivity, which may confound functional assays. In either of the previous scenarios, using primary tissue as a target cell source carries further issues since the ideal APC are typically either low frequency cell types that are difficult to culture (*e.g*. dendritic cells) or are cell types highly recalcitrant to genetic modification with viral vectors (*e.g*. B cells). For these reasons, a common strategy for producing large numbers of donor-derived APC for antigen screening is to create immortalized lines from donor cells by using laboratory strains of EBV to transform primary B cells into B- lymphoblastoid cell lines (56), which require long-term outgrowth for establishment and may display highly variable phenotypes between lines (57), or by stably expressing the proteins Bcl-6 and Bcl-xL in primary B cells (58,59) to produce long-lived germinal center-like B-cell lines, which is limited by the requirement for sustained presence of cytokine and CD40L-bearing feeder cells to maintain viability.

In addition to validating the basic YT-rCTL/KFRET synthetic chassis as a valuable tool for functional prototyping of recombinant TCR, we engineered the system to promote enhanced survival of K562-based sAPCs. Our primary motivation for this was to allow for reliable capture of cells marked for destruction by T cells by an early reporter of T cell cytotoxicity, before late-stage apoptosis and cellular self-destruction occur. We were able to preserve GZMB exposed cells until time points well beyond established peak signal time points. We also noted that KFRET cells, having been loaded with GZMB and undergoing FRET-shifting, could be separated from YT-rCTL effectors by FACS and successfully recovered and propagated in standard culture. Based on this, we suggest this system may be amenable to biopanning configurations in which functional screening of complex antigen libraries could be enhanced by iteratively co-culturing YT-rCTL with minigene library-bearing KFRET target cells, recovering FRET- shifted target cells by FACS, allowing target cells to recover and proliferate, and repeating the process multiple times before identifying response-eliciting peptides by minigene sequencing. Biopanning has been challenging for function-based assays of T cell response because target-centric response criteria have typically relied on complete APC destruction (60). In contrast, the YT-rCTL/KFRET system leverages early detection of effector cell attack on relevant target cells while protecting targets from destruction, presenting a possible solution to this challenge.

A current limitation of the YT-rCTL/KFRET system is the apparent lack of available markers for separating and isolating responder YT-rCTL clones from bulk co-culture to isolate effectors containing relevant TCR. In other experimental designs, this is commonly achieved by sorting on T cell activation induced markers or engineered reporters under the control of T cell specific transcription factors. These strategies enable screening of multiplexed libraries of TCR sequences against a desired antigenic target or group of targets and facilitate the selection of candidate TCR clonotypes from panels of pre-candidate TCR (45). Our preliminary RNA-seq survey of the YT- rCTL transcriptome did not identify activation induced markers, which points to a reporter system for detecting activation of individual YT-rCTL cells as an area for further development which, if successful, would expand the capability of the platform by allowing TCR library screening to be simultaneously coupled to the existing capability of the platform to perform antigen library screening.

## Conclusions

We have constructed and validated a recombinant system for characterizing TCR/pMHC functional responses. The YT-rCTL/KFRET cell chassis streamlines TCR assay workflows, from gene sequence to functional data, by eliminating requirements for sourcing and culturing experiment-specific primary tissue or cell lines. The GZMB FRET-shift functional read-out is considerably less time- and labor-intensive than classical T cell functional assays while being compatible with target-centric HTS configurations. We expect this technology will facilitate mapping TCR/pMHC interactions and characterizing clinical TCR candidates at the earliest stages of therapeutic development.

## Methods

### Plasmid DNA propagation and isolation

NEB Stable Competent E. coli was used for the propagation of DNA. Bacterial transformations were performed according to manufacturer protocol. E. coli was grown in LB broth at 30°C, shaking at 250 rpm. For solid medium, LB broth was supplemented with Bacto agar (1.5% [w/v]; Difco). Media were further supplemented with 100 μg/mL carbenicillin or 50 μg/mL kanamycin as appropriate. Plasmid DNA was isolated using Invitrogen PureLink HiPure Plasmid Filter Maxiprep Kit. All synthesized DNA was produced by Integrated DNA Technologies unless otherwise specified. All cloning enzymes were sourced from New England Biolabs unless otherwise specified. Sanger sequencing verification was performed at all intermediate steps by Genewiz.

### Construction of minigene and HLA DNA vectors

The lentiviral transfer plasmid was derived from the commercially available pCCL-c-MNDU3-PGK-EGFP backbone. To produce an acceptor cassette for minigene sequences or HLA allele sequences, the PGK-EGFP portion of the plasmid was replaced with custom synthesized cassettes to yield pMinigene-P2A-FRET, pMinigene-IRES-mStrawberry, pHLAI-P2A-mStrawberry, and pHLAI- P2A-FRET backbones by endonuclease cloning between the PacI/BamHI restriction sites in the original plasmid (BamHI site ablated during this step). Minigenes were synthesized and inserted into designated minigene positions via I-SceI/PI-SceI restriction endonuclease cloning. Coding sequences for HLA alleles of interest were either synthesized (for HLA-A*01:01) or sourced from the Riken DNA Bank (for HLA-C*14:03) and cloned into designated HLAI positions via NheI/MluI restriction cloning.

### Construction of TCR, CD3, and CD8 DNA vectors

The lentiviral transfer plasmid was derived from the commercially available pCCL-c-MNDU3-PGK-EGFP backbone. To produce an acceptor cassette for TCR, CD3, or CD8 sequences, the PGK-EGFP portion of the plasmid was replaced with a custom synthesized cassette to yield a pMND-MCS-P2A-mStrawberry backbone by endonuclease cloning between the PacI/BamHI restriction sites in the original plasmid (BamHI site ablated during this step). CD8 and CD3 cassettes were synthesized and cloned into pMND-MCS-P2A-mStrawberry via PacI/AscI restriction endonucleases to yield pCD8α-P2A-CD8β and pCD3δ- E2A-CD3γ-T2A-CD3ε-P2A-CD3ζ. EB81.103 and a3a TCR cassettes were synthesized as a single TCRɑ-T2A- TCRβ cloned into pMND-MCS-P2A-mStrawberry via BamHI/EcoRI restriction sites. HSDL1 mutant-reactive TCR was recovered from original T cell clone via 5’-RACE as individual ɑ- and β-chains (61), tailed with complete *TRAC* and *TRBC1* gene segments amplified from human genomic DNA (Promega), respectively, using overlap extension PCR, and sequentially cloned into pMND-MCS-P2A-mStrawberry via BamHI/NheI and MluI/EcoRI restriction sites, respectively.

### Construction of survival cassette and miRNA DNA vectors

The lentiviral transfer plasmid was derived from the commercially available pCCL-c-MNDU3-PGK-EGFP backbone. To produce a Golden Gate enabled destination vector, a synthetic DNA fragment comprising αLacZ and a Golden Gate cassette was inserted into ClaI/KpnI-linearized pCCL-c-MNDU3-PGK-EGFP using Gibson assembly to yield pMLV1-Destination plasmid. Golden Gate parts encoding survival cassette or perforin miRNA components were designed *in silico* to remove Type IIs restriction sites and any naturally occurring predicted caspase and granzyme cleavage sites, as well as predicted ubiquitination sites, and synthesized (Twist Biosciences). The pSurvival and pMIR series of plasmids were then created by combining the appropriate synthesized DNA parts with pMLV1-Destination in PaqCI-based Golden Gate assembly reactions.

### Mammalian cell culture

All cell cultures were maintained in RPMI-1640 supplemented with 2 mM GlutaMAX, 1 mM sodium pyruvate, 50 μM β-mercaptoethanol, 10 mM HEPES, 100 U/mL penicillin, 100 U/mL streptomycin, 1X MycoZap™ Prophylactic (Lonza) and 10% heat-inactivated fetal bovine serum. Culture media and supplements were all sourced from Gibco unless otherwise indicated. K562 and HEK-293T cells were sourced from ATCC. 721.221 cells were a kind gift from the Judy Liberman lab (Boston Children’s Hospital). YT-Indy cells were received under MTA from Indiana University. All cell cultures were maintained at 37°C and 5% CO_2_ atmosphere. K562 based cell lines were subcultured every 4 days by diluting cells 1/20. YT-Indy based cell lines were subcultured every 3 days by completely removing old media, washing 1X with PBS (Gibco), and re-seeding cultures at 1x10^6^ cells/well in 24-well gas permeable rapid cell expansion plate format (G-Rex, Wilson Wolf) . HEK-293T were passaged every 4 days by trypsinizing, washing 1X with PBS, and re-seeding cells at 4x10^6^ cells/flask in T-75 format. Cell counts were performed using an EVE automated cell counter. All cell lines were certified to be free of mycoplasma contamination using the Venor GeM Mycoplasma Detection Kit (Sigma) prior to experimental data collection.

### Primary T cell stimulation and expansion

Leukapheresis product (leukopaks) from healthy human donors were obtained from Stemcell Technologies shipped on wet ice. On receipt, leukopaks were processed using a Ficoll-Hypaque density gradient centrifugation and ACK buffer red-blood cell lysis procedure, both according to manufacturer provided protocols. The resulting isolated PBMCs were aliquoted into individual vials of 1x10^7^ cells and cryopreserved in liquid nitrogen vapor phase in heat-inactivated fetal bovine serum + 10% DMSO. Prior to activation, thawed PBMCs were FACS-sorted to isolate CD8^+^ CD4^-^ CD56^-^ cells (using BV785-conjugated anti-CD8 antibody, clone SK1 (Biolegend); PerCP-Cy5.5-conjugated anti-CD4 antibody, clone RPA-T4 (Biolegend); and FITC-conjugated anti-CD56 antibody, clone HCD56 (Stemcell)) and rested for 48 hours in complete RPMI + 300 IU/mL human IL-2 (Stemcell) at a cell density of 10x10^6^ cells/mL. T-cell activation was performed by transferring CD8^+^ T cells to wells of a non-TC treated, flat-bottom 96-well plate pre-coated with LEAF-purified anti-CD3 antibody, clone OKT3, and anti-CD28 antibody, clone 28.2 (eBioscience). Cells were stimulated for 20-24 hours on coated wells at 37°C and 5% CO2 atmosphere. After the stimulation period, cells were removed from coated wells, diluted to 1x10^4^ cells/mL in complete RPMI + 300 IU/mL hIL-2, and plated in U-bottom 96-well plates. Media was 50% changed every 4 days and used in co-culture experiments 11-14 days after stimulation.

### Virus production

Lentiviral vectors were produced by combining 180 µg of each transfer plasmid with 162 µg of pCMV-ΔR8.91 and 18 µg of pCMV-VSV-G plasmids. These DNA mixes were incubated with 18 mL OptiMEM and 1 mL of TransIT-LT1 reagent (Mirus) for 30 minutes at room temperature. To 12 x T-75 culture flasks containing between 16 – 24 million HEK-293T cells, old media was removed and replaced with 10 mL fresh, pre-warmed media and 1.5 mL of transfection mix per plate. Media was again replaced 18 hours post-transfection with 10 mL of pre-warmed fresh media. Viral supernatant was then collected at 48 hours post transfection (replacing with 10 mL pre-warmed fresh media) and 72 hours post-transfection. To concentrate virus, pooled supernatants were ultracentrifuged (100,000 RCF, 90 minutes, 4°C). Viral pellets were resuspended at 4°C overnight in 1 mL OptiMEM. Titers of viruses were determined by testing 2, 4, 8, 16, 32 or 64 μL of 10X diluted virus on 1x10^5^ K562 cell/well in a final volume of 500 μL of media in 24-well format. Transduction efficiency was determined by measuring the % of fluorescent cells (generated from encoded fluorescent protein or from antibody-staining transduction marker, as applicable) detected in flow cytometry. Values were used to determine K562-infecting units (KIU) per µL of undiluted virus concentrate.

### Creation of sAPC lines

K562 cell lines were transduced with viral vector produced using desired pHLAI-P2A- mStrawberry or pHLAI-P2A-FRET format constructs at an MOI of 1 KIU/cell. Purified HLA-expressing K562 were isolated by FACS to recover cells positive for RFP or FRET expression, respectively, and HLA expression (based on staining with APC-conjugated anti-human MHC, clone G46-2.6, BD Bioscience). Minigene-expressing target cells were produced by subsequently transducing HLA-expressing K562 with viral vector produced using pMinigene-P2A-FRET or pMinigene-IRES-mStrawberry format constructs at an MOI of 1 KIU/cell. Purified KFRET.HLA.minigene cell lines were isolated by FACS to recover cells positive for FRET or RFP expression, respectively. For survival cassette initial testing, KFRET.A0101.MAGEA3^160-183^ were additionally transduced with viral vector produced using pSurvival series of plasmids at an MOI of 1 KIU/cell. Purified survival cassette- expressing cell lines were isolated by FACS to recover cells positive for survival cassette transduction marker, truncated inert CD34 (using AF647-conjugated mAb clone 561, Biolegend).

### Creation of CRISPR-edited sAPC cells

The endogenous MAGEA3 epitope of EB81.103 TCR was excised from KFRET.A0101 cells by CRISPR using a two-guide approach. Epitope-flanking sgRNAs were designed with the Synthego CRISPR Design Tool. Cas9 and sgRNAs were purchased from Synthego. CRISPR editing was conducted by transfecting KFRET.A0101 cells with 10 pmol of Cas9 complexed to sgRNAs, at a 9:1 molar ratio, using a Neon Transfection System (Thermo) according to manufacturer-recommended K562-specific protocol. Transfected cells were recovered for 72 hours and single-cell cloned by FACS. To genotype cells, samples of outgrown clones were subject to gDNA isolation and PCR amplification of the edited locus using the primers 5’- GGATTTCCCAATTTGAG-3’ and 5’-GCTGACCTGGAGGAC-3’. The resulting amplicon was Sanger sequenced using a nested forward sequencing primer (Genewiz).

### Creation of rCTL lines

YT-Indy based rCTL effector cell lines were created by first double-infecting unmodified YT-Indy cell lines with viral vectors produced using pMND-CD3 and pMND-CD8 plasmids at an MOI of 0.05 KIU/cell of each to yield a CD8ɑβ^+^, CD3δɣεζ^+^ TCRɑβ^-^ YT-Indy “TCR-empty” cell line. Unsorted double-infected TCR-empty cells were subsequently transduced with viral vector produced using pMND-TCR series plasmid-of-interest at an MOI of 0.05 KIU/cell. Purified YT-rCTL.TCR cell lines were isolated by FACS to recover cells positive for CD3, CD8, and TCR (based on anti-CD3 and anti-CD8 antibody staining, and RFP expression). Primary TCR-T cells were created by transducing CD8^+^ T cells in conjunction with anti-CD3/28 stimulation (described above) with viral vector produced using pMND-TCR plasmid at an MOI of 30 KIU/cell. Purified TCR-T cells were isolated by FACS to recover cells positive for recombinant TCR (anti-TCR staining and RFP expression). For PRF miRNA initial testing, YT-rCTL.a3a were additionally transduced with viral vector produced using pMIR series plasmids at an MOI of 0.05 KIU/cell. Purified PRF miRNA cassette-expressing cell lines were isolated by FACS to recover cells positive for the transduction marker, truncated inert CD34 (using AF64-conjugated mAb clone 561, Biolegend).

### Granzyme FRET-shift assay co-cultures

To prepare assay co-cultures, Target and Effector cells were each adjusted to a density of 2 x 10^6^ cells/mL in fresh, pre-warmed media and 100 μL of each were separately added to 2 individual wells of a 96-well U-bottom plate. To initiate co-culture, one pair of effector/target wells was mixed and re-distributed; the other well pair was left unmixed and combined immediately before flow cytometry to serve as a T_0_ loading control. Co-culture plate was placed in 37°C and 5% CO2 atmosphere incubator for designated co-culture duration. On conclusion of co-cultures, cells were stained on-plate with 0.5% BV785-conjugated anti-CD8 antibody, clone SK1 (Biolegend) for 15 minutes at 4°C before harvesting co-cultures, diluting in 5X volumes of cold PBS, centrifuging at 300 x g for 5 minutes, and resuspending in 300 μL of cold PBS + 0.25% DRAQ7 (Thermo) per sample. Prepared co-culture samples were kept on ice and taken immediately for flow cytometry/FACS. Co-cultures were scaled for some experiments by preparing more U-bottom wells as needed (each well consisting of 1 x 10^5^ Targets and 1 x 10^5^ Effectors per well).

### Flow cytometry/FACS

All flow cytometric analysis was performed on BD LSRII Fortessa and all cell sorting was performed on BD FACSAria Fusion. Relative cell number quantitation was performed by holding flow rate and acquisition time constant (along with resuspension volume in sample preparation). Gating was performed by monitoring Fixable Viability Dye eFluor 780 channel (ex. 640, em. 780/60 + 750LP), DRAQ7 channel (ex. 640, em. 730/45 + 690LP), RFP channel (ex. 561, em 610/20 + 600LP), YFP channel (ex. 488, em. 530/30 + 505LP), APC/AF647 channel (ex. 640/em. 670/14), BV785 channel (ex. 405, em. 780/60 + 750LP), FRET channel (ex. 405, em. 525/50 + 505LP), and/or CFP channel (ex. 405, em. 450/50) according to experiment design. For GZMB and PRF intracellular staining assays, cells were fixed and permeabilized using Biolegend reagents and manufacturer protocols.

### RNA-seq

Samples of unmodified YT-Indy cells, YT-rCTL.a3a cells, antigen-experienced YT-rCTL.a3a cells FACS-isolated from 12-hour co-culture with KFRET.A0101.MAGEA3^160-183^, primary CD8^+^ a3a TCR-T cells, and antigen-experienced primary CD8^+^ a3a TCR-T cells FACS-isolated from 12-hour co-culture with KFRET.A0101.MAGEA3^160-183^ (1 x 10^6^ cells of each) were pelleted, flash-frozen, and shipped on dry-ice for RNA-seq (Genewiz). Primary a3a TCR-T cells used in this study were prepared as described above from bulk frozen donor PBMC and were sorted/cryopreserved for sequencing 9 days after anti-CD3/28 stimulation. Illumina library construction was conducted on polyA-isolated RNA and prepared libraries were subject to PE150 sequencing using the Illumina HiSeq platform. Reads were aligned to the reference human transcriptome using STAR and analyzed for differential gene expression using DESeq2.

## Declarations

### Ethics approval and consent to participate

Commercially-sourced primary human tissue used in this study was obtained from suppliers using Institutional Review Board- or Research Ethics Committee-approved consent forms and protocols.

### Availability of data and materials

Raw data generated in this study are available from the lead or corresponding author on reasonable request. Raw .fastq files from RNA-seq study are available in the Zenodo repository 10.5281/zenodo.8428844.

### Author contributions

GS ideated the technology and experimental strategy with guidance from RAH. GS, ZA, and JR designed and prepared all DNA constructs. GS, FT, CM, and EY prepared viral vectors and transduced cell lines. JR and FT conducted CRISPR editing and clonal selection. GS and FT performed all cellular assays. Data analysis and manuscript preparation was done by GS with editing and feedback from all authors.

## Acknowledgements

The authors would like to acknowledge Dr. Rodrigo Vasquez-Lombardi and Dr. Daniel Woodsworth for their valuable input and ideas.

---

**Supplementary Figure 1.**
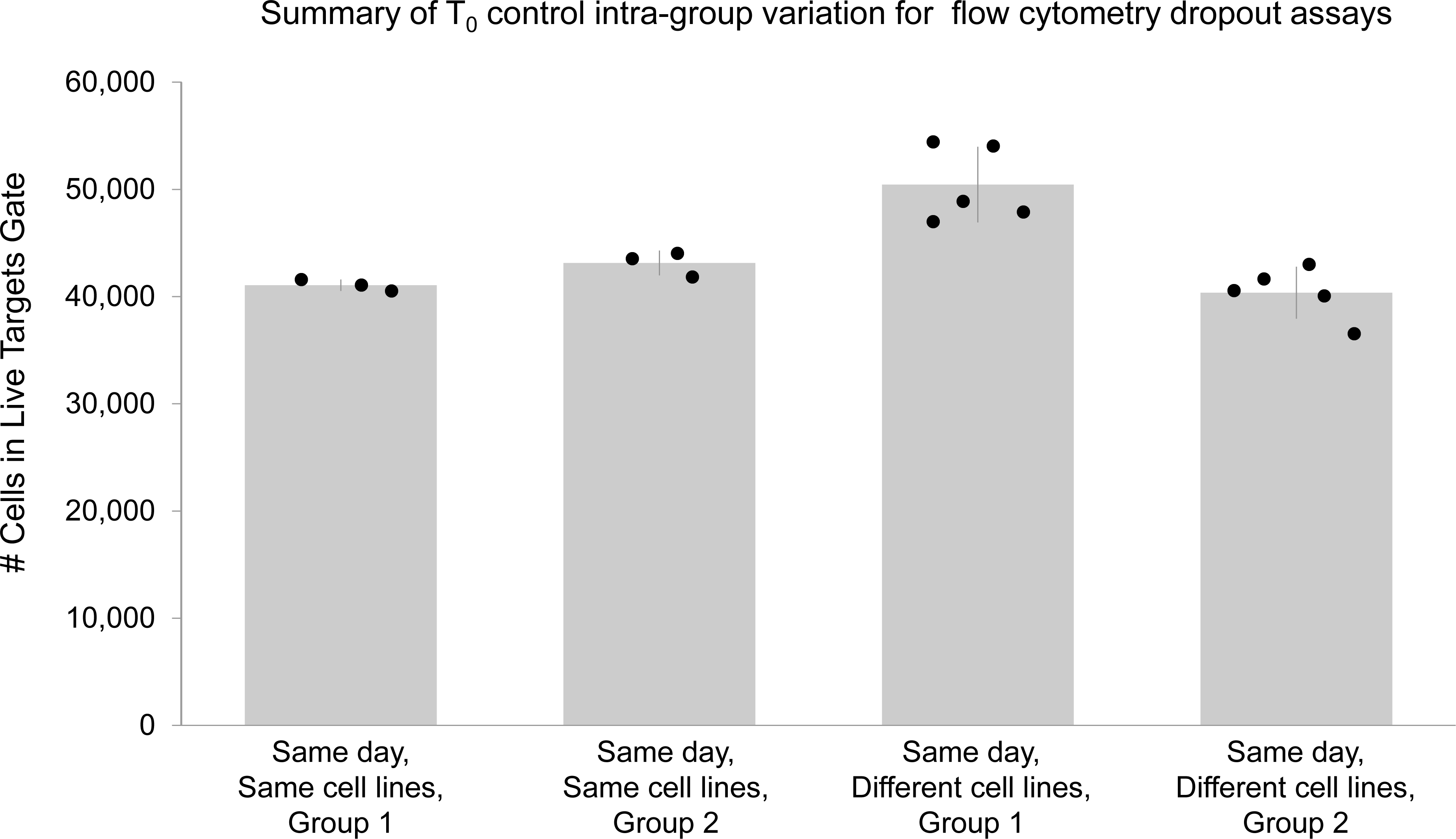
Intra-experiment variance of T_0_ controls is consistent. A summary of cell counts obtained in live targets gate of YT-rCTL/KFRET T_0_ controls illustrates variance between absolute counts in individual samples. While statistically significant differences were apparent between groups prepared for different experimental runs (*e.g.*, on different days), samples prepared in the same experimental run (*e.g.* on the same day) are found to be highly consistent, even when comparing T_0_ controls composed of differing cell lines. By Levene’s test, the variance between groups of samples prepared in different batches is found to be homogenous, indicating that the use of relative cell counting as a measure of cell dropout over co-culture duration is reliable when matched experimental and T_0_ samples are prepared in parallel.

**Supplementary Figure 2.**
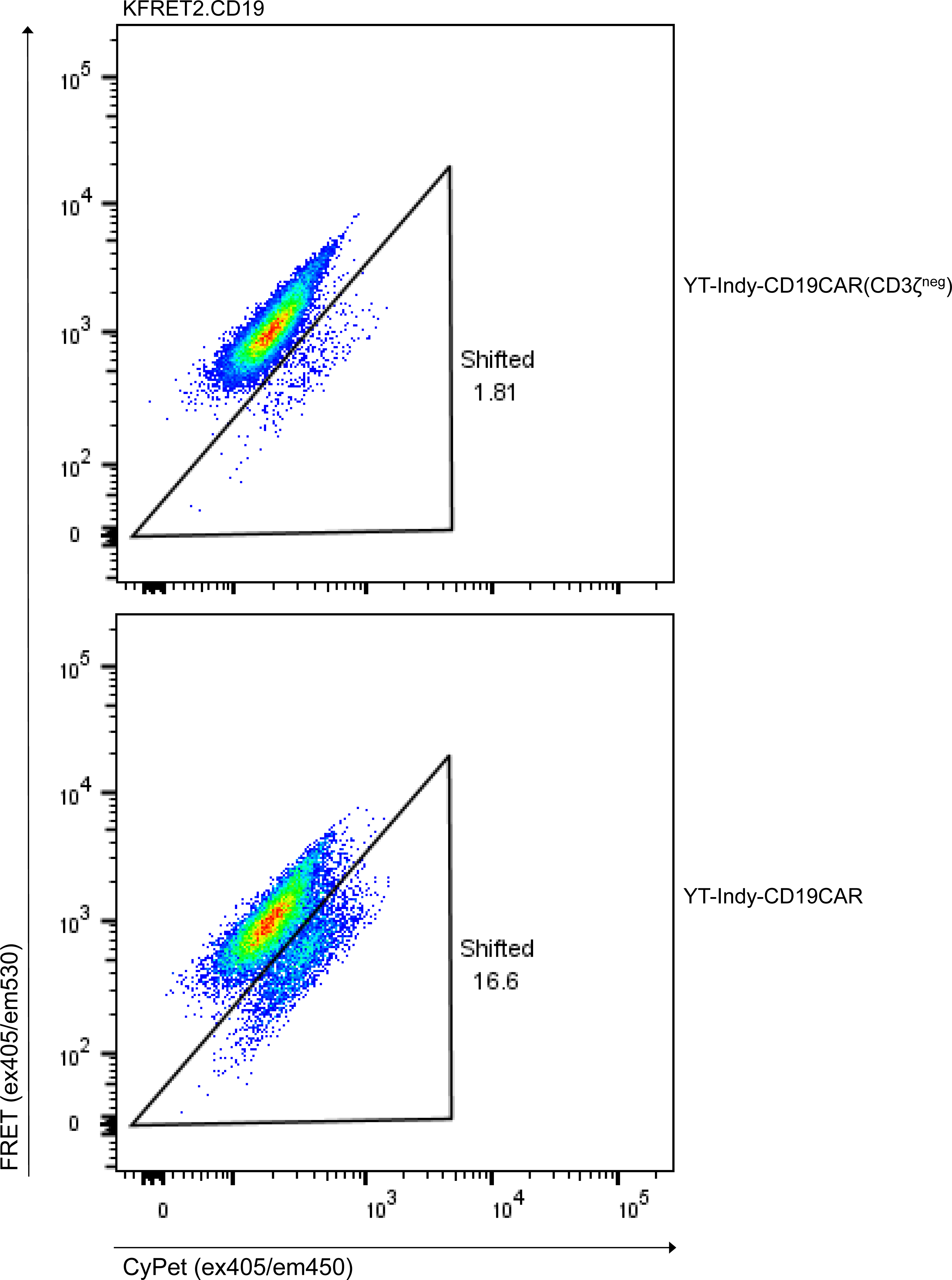
CAR-mediated cytotoxicity is primarily triggered through the CD3ζ signaling domain. K562 cells modified to express FRET2 reporter and CD19 protein coding sequence were co-incubated with YT-Indy modified with either an FMC63-41BB-CD3ζ anti-CD19 CAR or a prematurely truncated FMC63- 41BB anti-CD19 CAR (CD3ζ^neg^) at 1:1 effector:target ratios for 4 hours.

**Supplementary Figure 3.**
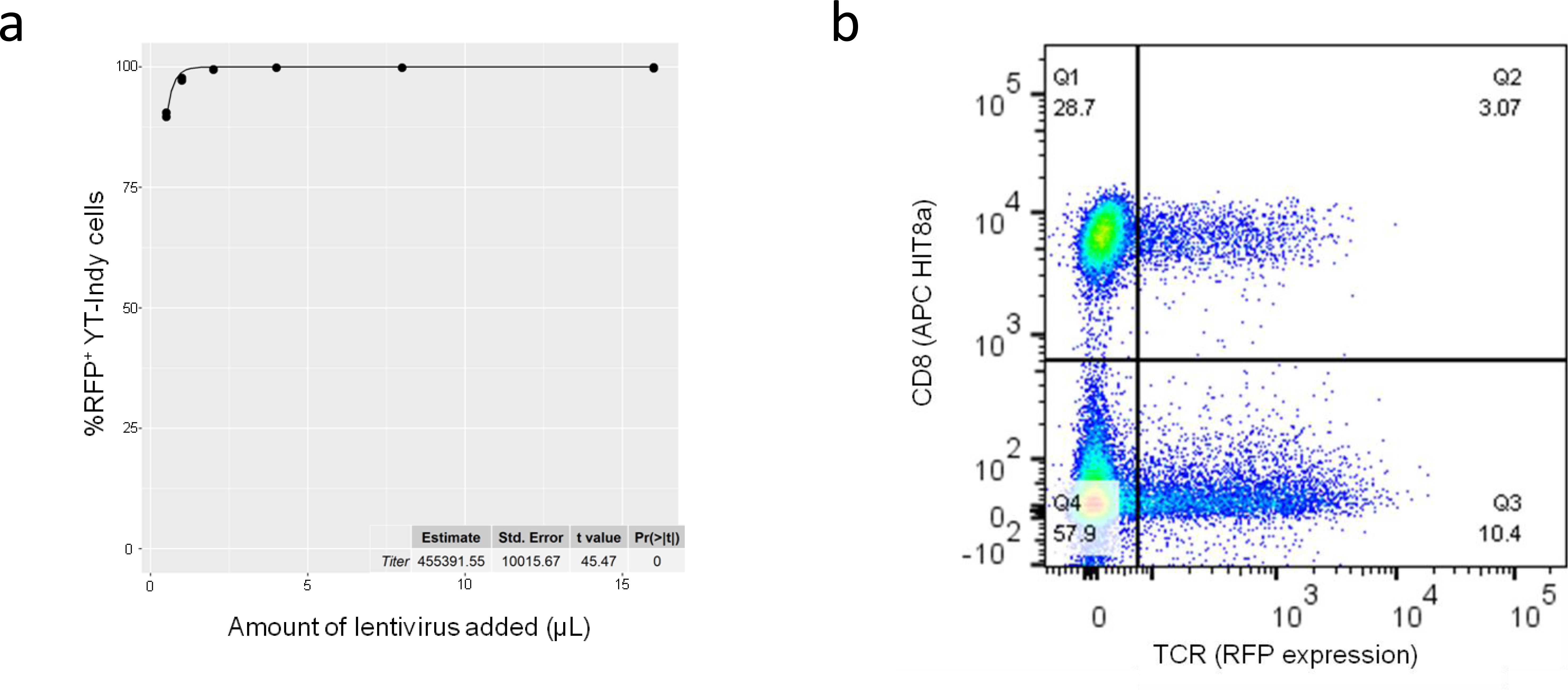
Lentiviral vector uptake is three orders of magnitude more efficient in YT-Indy cells than primary T cells. TCR-2A-RFP encoding lentiviral vector was functionally titered over YT-Indy cells by adding increasing amounts of virus to a fixed number of cells in a fixed volume. The resulting fluorescence was measured by flow cytometry 72 hours later. The proportions of positive cells in each condition were fit to a Poisson probability mass function by nonlinear least squares regression to determine a functional titer of 4.55 x 10^5^ infectious units/μL against YT-Indy (a). The same virus was also applied to an activated and proliferating culture of human T cells. In this experiment 150 μL of viral vector was added to 5x10^5^ T cells to yield a transduction efficiency of 13.47% (b). Applying Poisson probability to this result and accounting for input cell and virus quantities, the same product had an effective functional titer of 483 infectious units/μL against T cells.

**Supplementary Figure 4.**
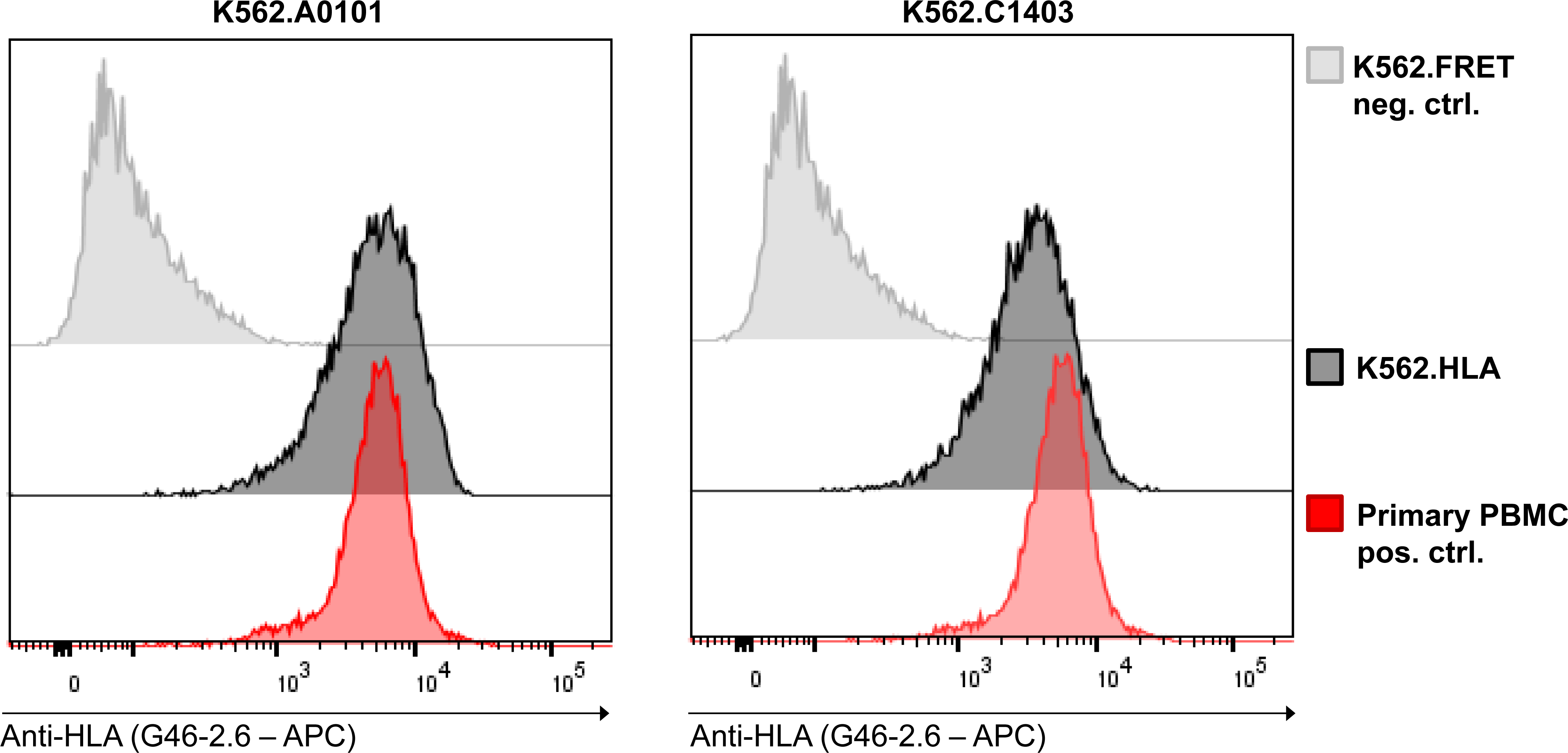
Robust recombinant expression of MHC molecules on K562 cells following transduction of HLA allele coding sequences. K562 cells transduced with lentivirus at an MOI of 1 KIU/cell encoding either HLA-A*01:01 or HLA-C*14:03 allele coding sequences were purity-sorted by FACS to isolate RFP and surface MHC-expressing cells. Recovered cell lines were characterized with respect to surface expression level, as measured by MFI from staining and flow cytometry using pan-MHC antibody, relative to primary human PBMC positive control.

**Supplementary Figure 5.**
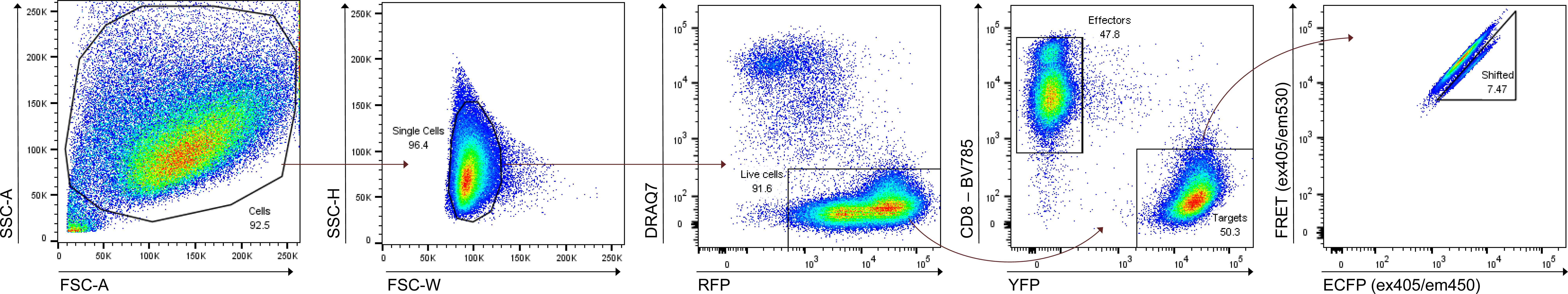
Representative gating strategy for FRET-shift flow cytometry in YT-rCTL/KFRET experiments. A common flow cytometry template was employed to facilitate endpoint measurements of all YT- rCTL/KFRET format TCR/pMHC functional assays shown in Main Figures 3, 4, and 5, and Supplementary Figures 5, 6, 7, 8, and 9. Intact, single cells are first selected using sequential forward- and side-scatter gates. Viable cells are then selected by gating on cells negative for DRAQ7 vital staining dye and positive for RFP living color. Effector and target cells are separated by CD8-BV785 and EYFP expression. Unshifted and Shifted gates are drawn by adjusting gate boundaries to keep background FRET-shift in T_0_ or targets-only control samples to <1%.

**Supplementary Figure 6.**
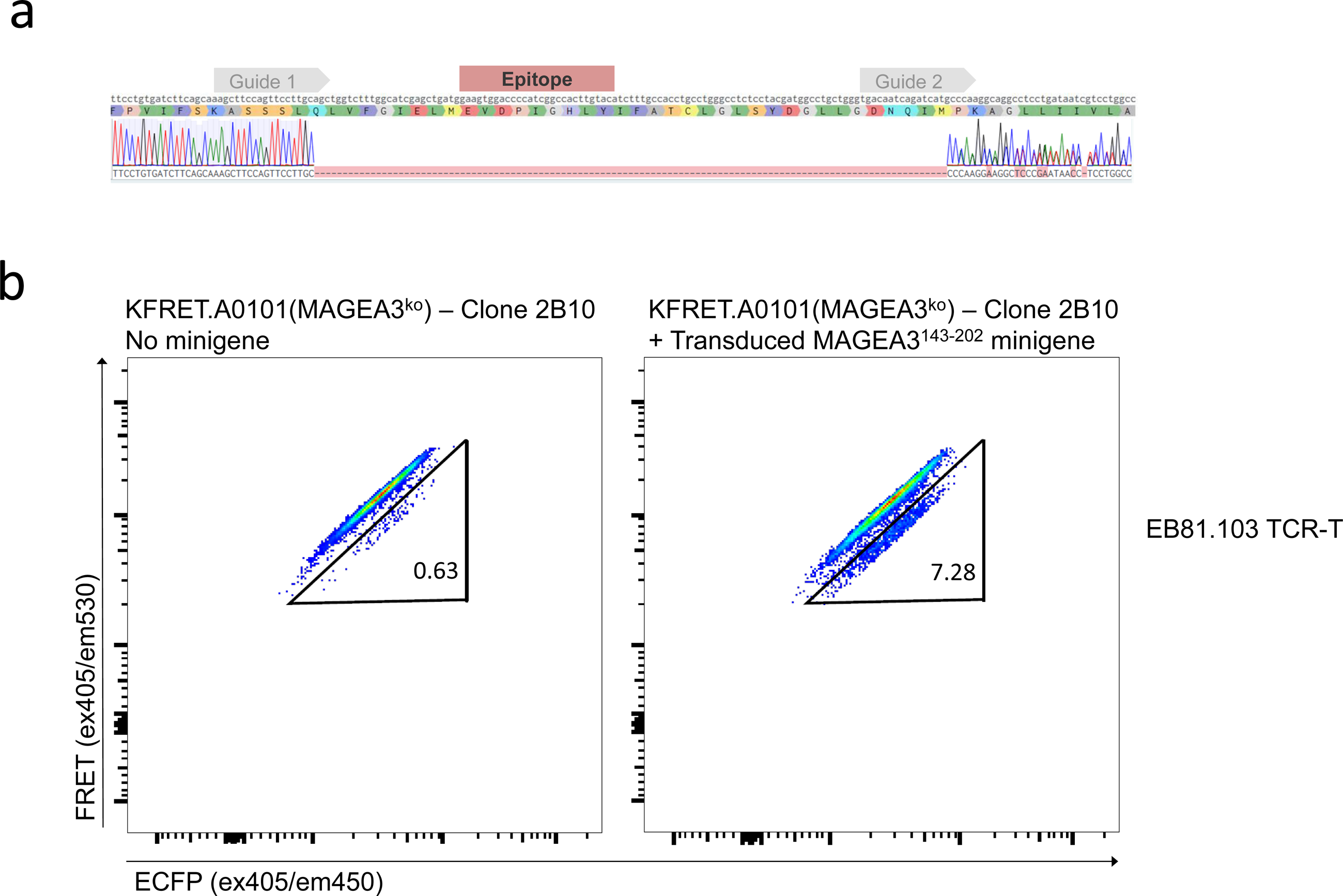
Successful excision of endogenous MAGEA3 epitope of EB81.103 TCR from KFRET.A0101 cells. Upon electroporation with epitope-flanking sgRNAs in complex with Cas9 protein, KFRET.A0101 cells were single-cell sorted to yield 384 founder clones for selection. Colonies with successful outgrowth were individually checked for sustained RFP and FRET expression by flow cytometry. Phenotypically selected cell lines were then genotyped by PCR amplification and Sanger sequencing of the genomic region around the edited locus. (a) The clone ultimately selected for further study (2B10) showed an unambiguous deletion of the desired site in Sanger trace. The presence of multiple traces downstream of the edited site indicates that multiple distinct successful edits were made. Since K562 has maintained X-chromosome ploidy of 2, ablation of the MAGEA3 epitope was assumed to be complete. (b) The selected 2B10 clone was transduced with MAGEA3^143-202^ minigene to restore the deleted epitope. A rescue experiment in which primary EB81.103 TCR-T cells were co-cultured with either KFRET.A0101(2B10) or KFRET.A0101(2B10).MAGEA3^143-202^ at 1.5:1 effector:target ratio (to account for ∼60% TCR transduction efficiency in T cells) for 12 hours demonstrates deletion of endogenous MAGEA3 epitope and maintained function of the HLA-A*01:01 and GZMB FRET-reporter transgenes in the 2B10 clone.

**Supplementary Figure 7.**
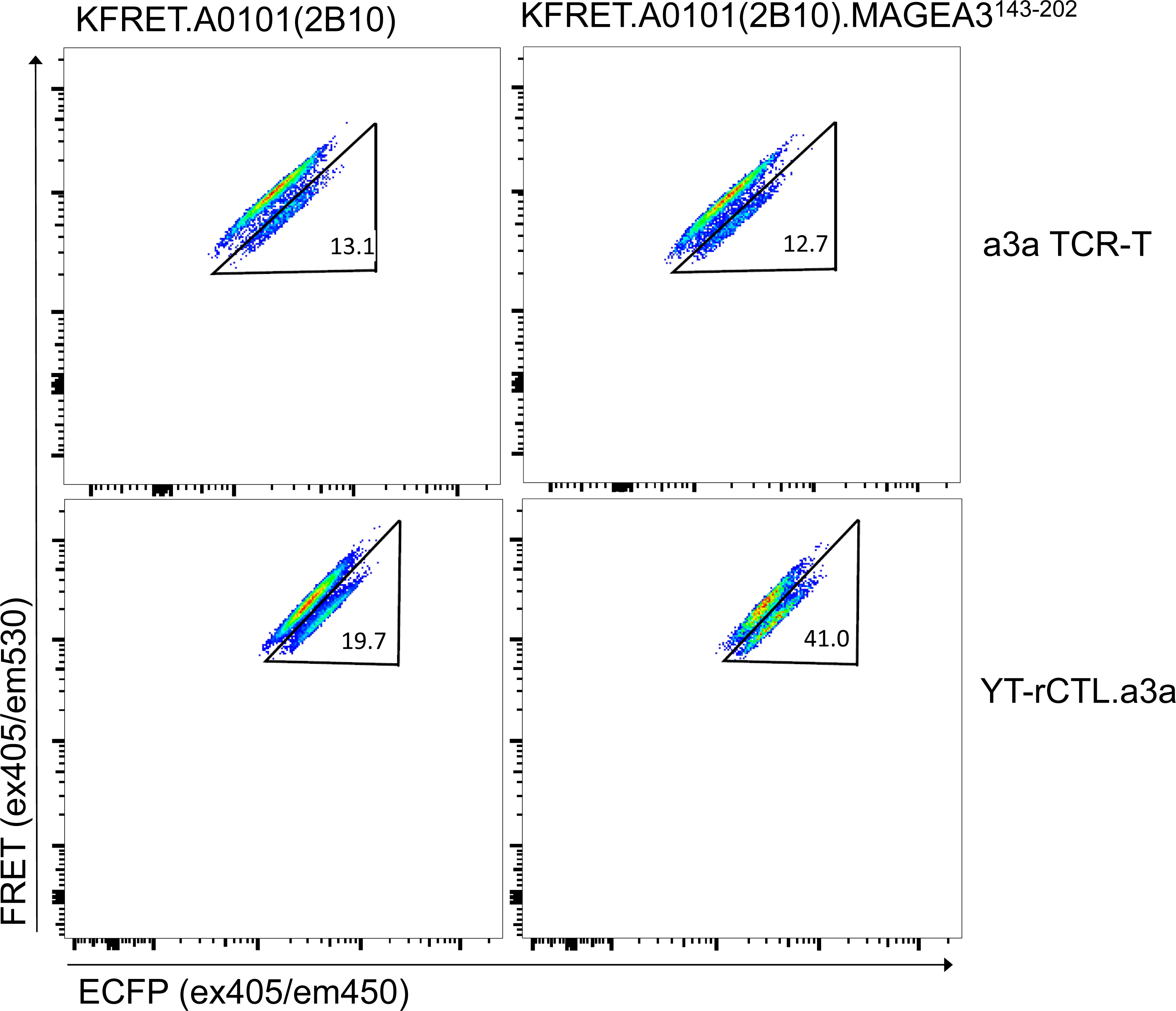
The a3a variant of EB81.103 results in a larger percent FRET-shift signal than wild-type. KFRET.A0101(2B10) and KFRET.A0101(2B10).MAGEA3^143-202^ target cells were co-cultured with either YT-rCTL.a3a or primary cytotoxic T cells freshly isolated by FACS purification CD8^+^ CD4^-^ CD56^-^ T cells from healthy donor PBMC, activated by plate-bound anti-CD3/28 stimulation, and transduced with a3a TCR lentivirus. YT-rCTL.a3a were co-cultured with targets at a 1:1 effector:target ratio for 12 hours. Primary a3a TCR-T cells were co-cultured at a 1.5:1 ratio (to compensate for 68% TCR transduction efficiency) for 12 hours.

**Supplementary Figure 8.**
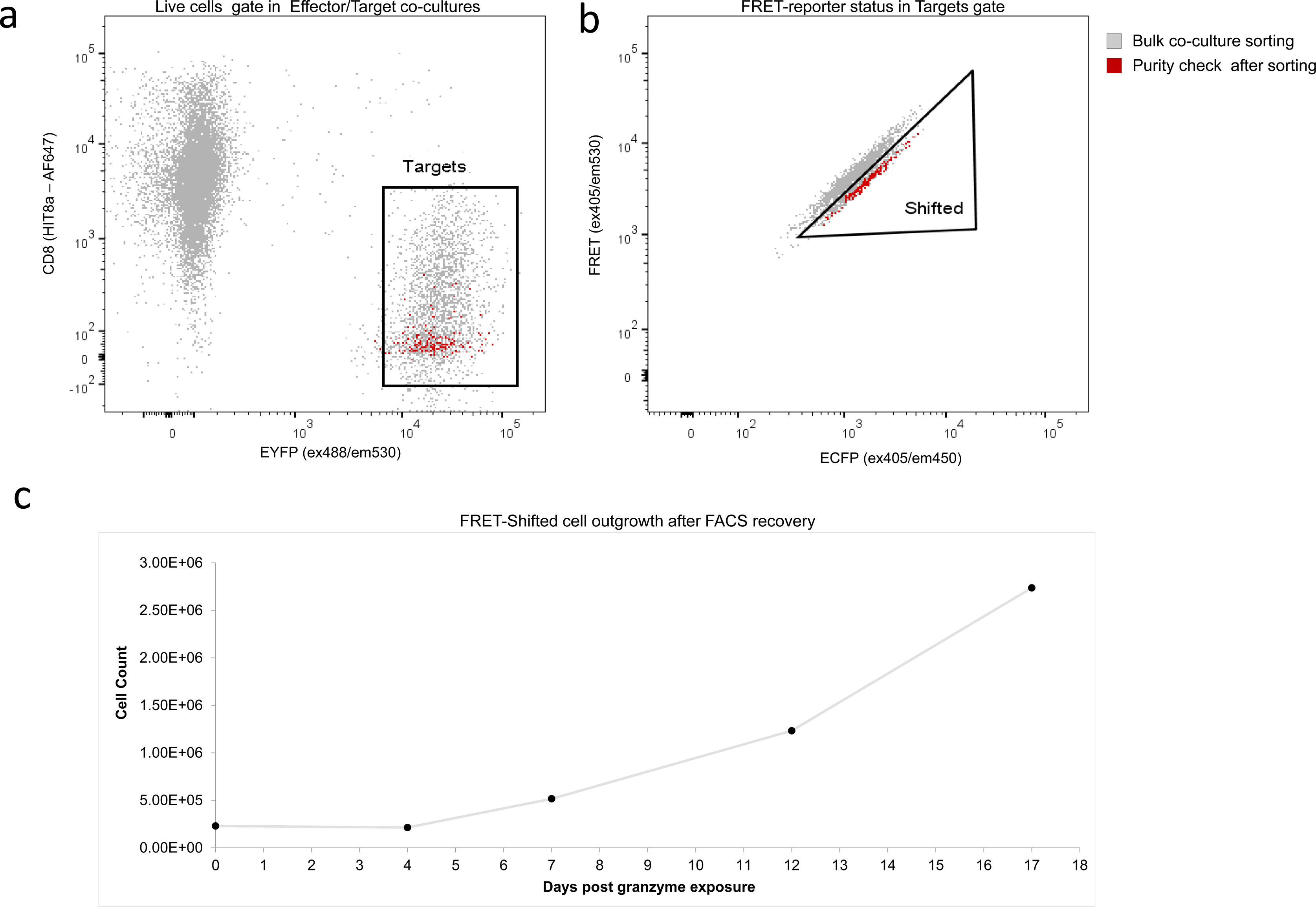
Outgrowth of FRET-shifted cells after their recovery by FACS. YT-rCTL.a3a and KFRET.A0101.MAGEA3^164-179^ peptide were co-cultured at a 1:1 effector:target ratio for 12 hours. Target sAPC cells undergoing FRET-shift were isolated by FACS. Recovered cells were immediately assessed for purity by flow cytometry and were determined to be 100% free of any contaminating effector cells (a) or contaminating non-FRET- shifted target cells (b). Recovered FRET-shifted cells (3.25 x 10^5^ total) were placed into fresh culture media and counted at regular intervals using an automated trypan based cell counter. Cell counts initially declined and reached a minimum total cell count of 2.14 x 10^5^ by day 4, but subsequently reversed their decline and followed an exponential growth curve post-day 4 (c).

**Supplementary Figure 9.**
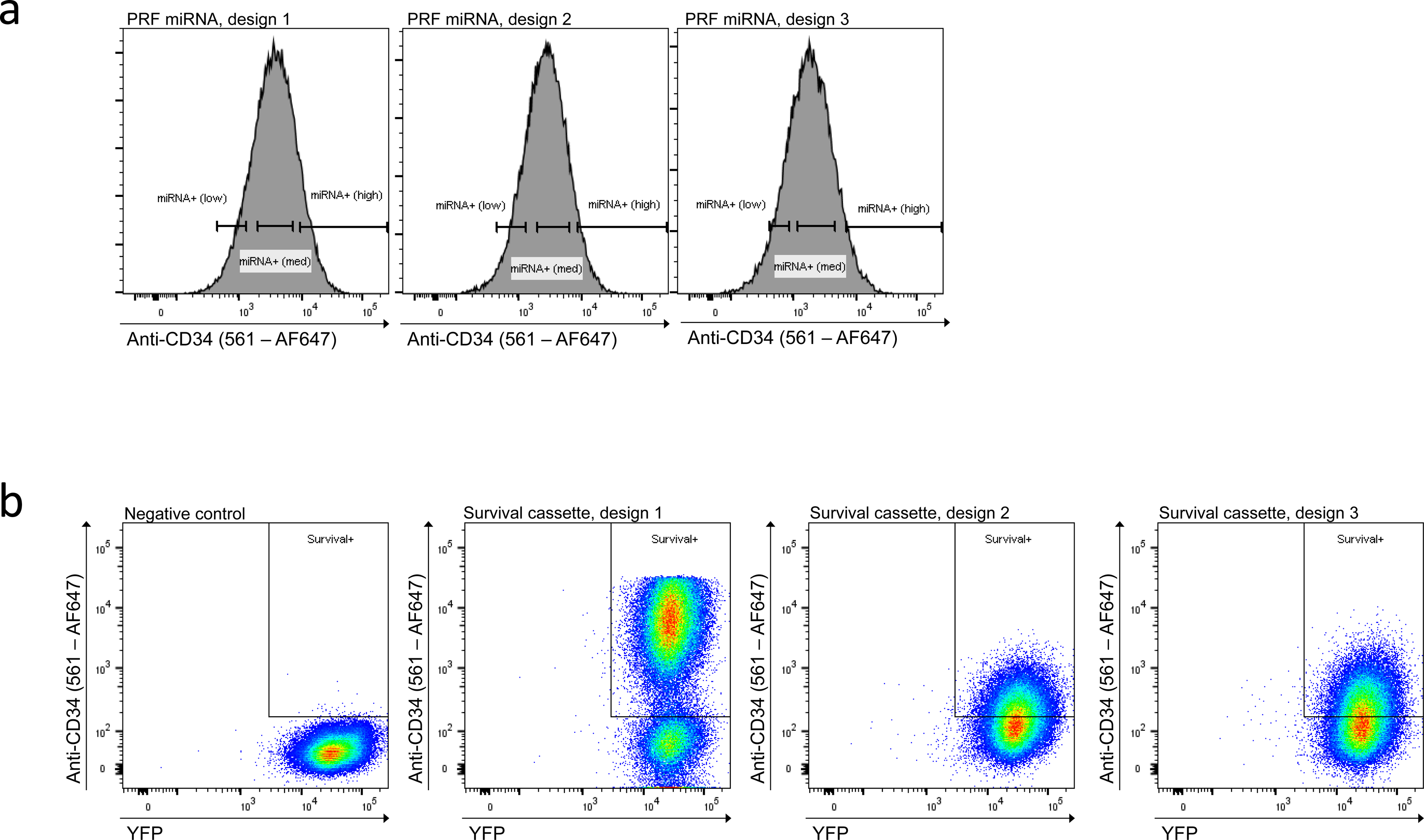
FACS purity-sorting schema for KFRET Survival cassette lines and YT-rCTL PRF miRNA lines. (a) KFRET.A0101.MAGEA3^160-183^ cells were transduced at an MOI of 1 KIU/cell with each of three Survival cassette designs or an empty vector control. Purity-sorting was conducted 72 hours after transduction by staining cells with anti-CD34 antibody to detect inert truncated recombinant CD34 transduction marker, and gating on live target cells with AF647 MFI using a threshold of >5 standard deviation above the mean of the negative control. (b) YT-rCTL.a3a cells transduced at an MOI of 0.05 KIU/cell with each of three PRF miRNA designs or a scrambled miRNA control. Purity-sorting was conducted 72 hours after transduction by staining cells with anti- CD34 antibody to detect inert truncated recombinant CD34 transduction marker. Gating on AF647 signal was done by arbitrarily selecting low, medium, and high expression gates for each. The minimum threshold of the low expresser population was set at 5 standard deviations above the mean of the scrambled control.

**Supplementary Figure 10.**
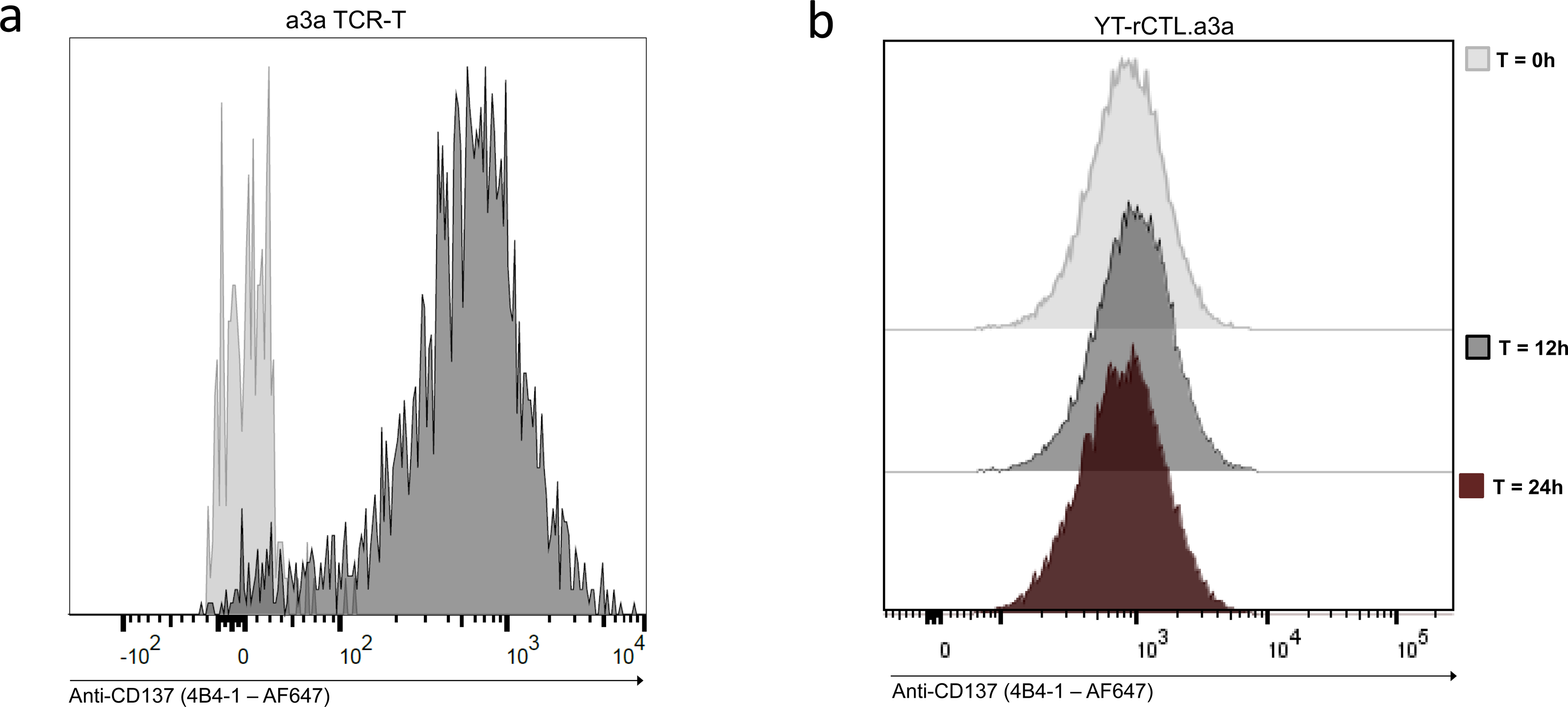
Assessing CD137 upregulation in YT-rCTL cells. (a) Primary CD8^+^ a3a TCR-T cells co-cultured with KFRET.A0101.MAGEA3^160-183^ sAPC were stained with anti-CD137 antibody (AF647-conjugated clone 4B4-1, Biolegend) and analyzed by flow cytometry. Clear resolution between T_0_ control and 12-hour co-culture experiment was observed. (b) YT-rCTL.a3a effectors co-cultured with KFRET.A0101.MAGEA3^160-183^ sAPC for either 12 or 24 hours were stained with anti-CD137 antibody and analyzed by flow cytometry. No resolution between T_0_ control and either the 12-hour or 24-hour co-culture experiments were observable.

